# Coupling of dynamic microtubules to F-actin by Fmn2 regulates chemotaxis of neuronal growth cones

**DOI:** 10.1101/2020.01.18.911131

**Authors:** Tanushree Kundu, Priyanka Dutta, Dhriti Nagar, Sankar Maiti, Aurnab Ghose

## Abstract

Dynamic co-regulation of the actin and microtubule subsystems enables the highly precise and adaptive remodeling of the cytoskeleton necessary for critical cellular processes, like axonal pathfinding. The modes and mediators of this interpolymer crosstalk, however, are inadequately understood.

We identify Fmn2, a non-diaphanous related formin associated with cognitive disabilities, as a novel regulator of cooperative actin-microtubule remodeling in growth cones. We show that Fmn2 stabilizes microtubules in the growth cones of cultured spinal neurons and also *in vivo*. Superresolution imaging revealed that Fmn2 facilitates guidance of exploratory microtubules along actin bundles into the chemosensory filopodia. Using live imaging, biochemistry and single-molecule assays we show that a C-terminal domain in Fmn2 is necessary for the dynamic association between microtubules and actin filaments. In the absence of the cross-bridging function of Fmn2, filopodial capture of microtubules is compromised resulting in de-stabilized filopodial protrusions and deficits in growth cone chemotaxis.

Our results uncover a critical function for Fmn2 in actin-microtubule crosstalk in neurons and demonstrate that modulating microtubule dynamics via associations with F-actin is central to directional motility.

**SIGNIFICANCE STATEMENT:** The formin family member, Fmn2, is associated with cognitive impairment and neurodegenerative conditions though its function in neurons is poorly characterized. We report a novel actin-microtubule cross-bridging activity for Fmn2 that facilitates efficient targeting and capture of microtubules in growth cone filopodia. This activity is necessary for accurate pathfinding of axons and may contribute to Fmn2-associated neuropathologies.

The precision and adaptability of cytoskeleton-driven processes are intimately dependent on the coupled activities of its component systems. Our study identifies a novel modality of co-regulated remodelling of the actin and microtubule cytoskeletons that facilitate critical cellular behaviour like neuronal chemotaxis.

## INTRODUCTION

The actin and microtubule subsystems of the cellular cytoskeleton are essential for the organization of the cytoplasm, cell and tissue morphogenesis, and cellular motility. In recent years, it is increasingly apparent that crosstalk between these two cytoskeleton systems is essential for regulating core cellular processes and their dynamic remodeling is functionally coordinated (Coles & Bradke, 2015, Dogterom & Koenderink, 2019). Growth cone -mediated axonal extension and pathfinding necessitates dynamic coordination between the peripheral filamentous actin (F-actin) and the exploratory microtubules (MTs) and is necessary for the establishment of neural circuits (Coles & Bradke, 2015, Dogterom & Koenderink, 2019, Lowery & Van Vactor, 2009, Vitriol & Zheng, 2012).

The central (C) domain of the growth cone is enriched with stable MTs extending from the axon shaft. While most of the stable MTs are confined to the C-domain, few dynamic MTs explore the peripheral (P)-region and interact with F-actin structures in the leading edge lamellipodia and filopodia. Filopodia are chemosensory structures composed of parallel F-actin filaments essential for chemotactic guidance.

Stabilization of filopodia, in the direction of growth cone turning, mediates directional motility of growth cones. Guidance cue signalling biases the activity of exploratory MTs to the direction of turning, increasing their entry and stabilization within filopodia on that side (Buck & Zheng, 2002, Mack et al., 2000, Sabry et al., 1991, Williamson et al., 1996). Biased MT stabilization in filopodia, by ensuring long-lived tracks for cargo transport and increasing mechanical resilience, supports filopodial stability and promotes growth cone turning. The entry and capture of MTs in filopodia involves dynamic associations of the exploratory MT and the F-actin structures in the transition (T) and P-domains of the growth cones. The F-actin bundles of filopodia appear to serve as tracks for extending MTs (Schaefer et al., 2002, Zhou et al., 2002) and may facilitate growth cone extension. The activities mediating the association of MTs with F-actin in neuronal growth cones are only beginning to be identified (Biswas & Kalil, 2018, Liu et al., 2010, Slater et al., 2019, Szikora et al., 2017).

Few proteins, especially some MT plus-end tracking proteins (+TIPs), have been identified to mediate aspects of actin-MT crosstalk however the precise mechanisms mediating actin-MT interactions are not well understood. In non-neuronal cells, another class of proteins, the formins, have also been implicated in actin-MT cross-talk. Formins are a family of 15 proteins (in mammals), most of which are also expressed in the nervous system (Dutta & Maiti, 2015, Krainer et al., 2013). These multi-domain proteins are commonly defined as actin nucleators but they are emerging as cytoskeleton regulators with a diversity of functions and spatiotemporal regulation. Recent work has identified the ability of some formins to regulate MT dynamics. The +TIP protein, CLIP-170 has been shown to associate with the formin, mDia1 and accelerate F-actin polymerization at MT plus ends (Henty-Ridilla et al., 2016). Other members of the Diaphanous-related formins (DRFs) have been found to influence MT stability in non-neuronal cells. Much less is known about formin-mediated actin-MT crosstalk in neurons. mDia1 activity has been associated with amyloid β-induced increase in MT stabilization in rodent neurons (Qu et al., 2017). In *Drosophila* neurons, the DRF formin DAAM has been shown to crosslink F-actin and MTs (Szikora et al., 2017). The expression of several formins in mammalian neurons suggests rich functional diversity of this family in the nervous system conferring functional precision and adaptability to the neuronal cytoskeleton.

Formin 2 (Fmn2) is a non-DRF formin whose expression is enriched in developing and mature mammalian nervous systems (Leader & Leder, 2000, Sahasrabudhe et al., 2016). Biallelic loss of Fmn2 and heterozygous deletions in Fmn2 has been associated with human intellectual disability (Almuqbil et al., 2013, Law et al., 2014). Mutations in Fmn2 have been found in a subpopulation of children with sensory processing dysfunction (Marco et al., 2018). Fmn2 expression was reduced in post-mortem brain samples of patients with post-traumatic stress disorder and Alzheimer’s disease (Agis-Balboa et al., 2017). In rodents, loss of Fmn2 has been associated with accelerated age-associated memory impairment and amyloid-induced deregulation of gene expression (Agis-Balboa et al., 2017, Peleg et al., 2010). However, little is known about the mechanistic underpinnings of Fmn2 function in the brain. We have previously shown that Fmn2 regulates axonal outgrowth and pathfinding in chick spinal commissural neurons *in vivo* (Sahasrabudhe et al., 2016). This study identified Fmn2 function in stabilising the adhesive contacts of the growth cone with the extracellular matrix (ECM) and consequently necessary for efficient growth cone translocation. Further, reduced Fmn2 in spinal commissural neurons displayed aberrant axonal trajectories *in vivo* and was suggestive of chemotactic deficits (Sahasrabudhe et al., 2016).

As deficits of chemotactic responses could result from aberrant MT dynamics, we explored Fmn2 function in regulating MT dynamics in growth cones. We discovered that Fmn2 facilitates the exploration of MTs into the P-domain of the growth cones by guiding MT into filopodia by physically coupling them to the F-actin bundles. This crosslinking is necessary to stabilize MTs. *In vitro* reconstitution assays confirm F-actin-MT cross-bridging by Fmn2 via its C-terminal FSI domain. Fmn2-mediated MT capture in filopodia appears to be critical for filopodial dynamics and stability and regulates growth cone turning. This work identifies Fmn2 as novel mediator of synchronised co-organization of the F-actin and MT cytoskeletons in neuronal growth cones underscoring Fmn2 function in neural development.

## RESULTS

### Fmn2 regulates microtubule organization in neuronal growth cones

Previously we have identified Fmn2 as a regulator of growth cone motility and guidance, both *in vitro* and *in vivo* (Sahasrabudhe et al., 2016). Fmn2 is localized to F-actin rich structures, like along the length of filopodial F-actin bundles in neuronal growth cones and stress fibres in fibroblasts (Sahasrabudhe et al., 2016). As MTs are involved in growth cone steering and are themselves subject to guidance along actin bundles (Schaefer et al., 2002), we investigated the role of Fmn2 in regulating MT dynamics in the growth cone.

Immunolocalization of Fmn2 in growth cones of chick spinal commissural neurons showed enrichment of Fmn2 along the length of filopodial F-actin bundles extending all the way to the base of these bundles (Figure 1A-A”’; also shown earlier in (Sahasrabudhe et al., 2016)). To test Fmn2 function in MT organization, we depleted Fmn2 using specific translation blocking morpholinos (Fmn2-MO; see Materials and Methods for characterization) that resulted in at least 75% reduction in endogenous Fmn2 levels (Figure S1A) followed by evaluation of MTs (anti α-tubulin immunofluorescence) and F-actin (phalloidin immunofluorescence) using STED nanoscopy (Figure 1 B,C). As reported earlier (Sahasrabudhe et al., 2016), depletion of Fmn2 reduced the number and length of growth cone filopodia compared to non-specific control morpholino (Ctl-MO)-treated neurons (Figure 1 D,E). Interestingly, Fmn2-MO-treatment resulted in a reduced proportion of filopodia with detectable MTs (Figure 1 F; 41.47 ±5.13% in Ctl-MO-treated neurons to 23.88 ±6.01% following Fmn2 knockdown). Further, though the number of MT ends in the growth cones were comparable between Ctl-MO and Fmn2-MO transfected neurons (Figure 1 G), the fraction of MTs invading filopodia was reduced by almost 50% upon Fmn2 depletion (Figure 1 H; 28.77 ±3.90% in control treatments to 13.66 ±3.69% in Fmn2-MO transfected growth cones).

**Figure 1:**
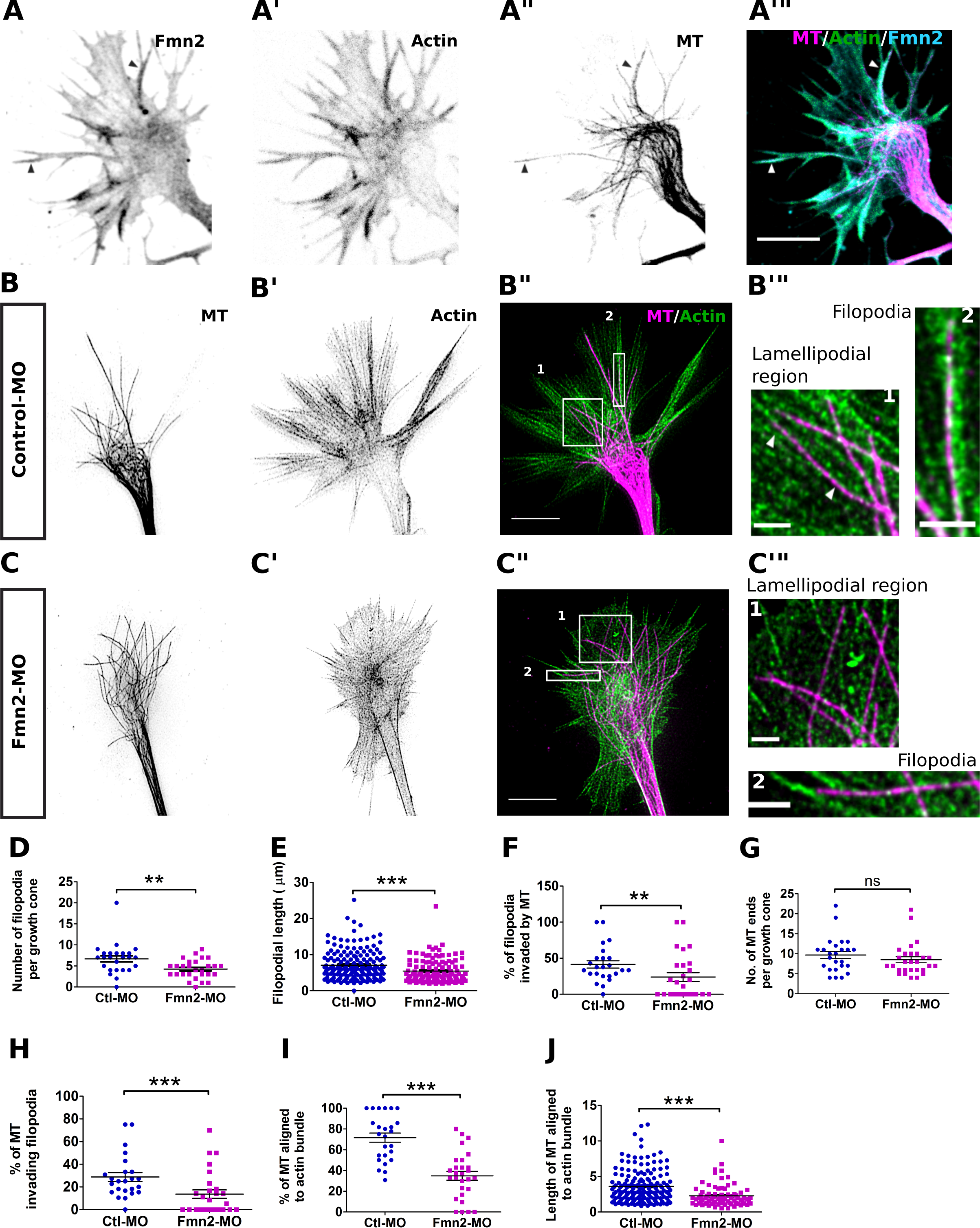
Fmn2 co-localizes with F-actin and microtubules (MT) in filopodia and promotes actin-MT alignment in the peripheral regions of the growth cone. (A-A’’’) Representative micrographs of triple immunostaining using specific antibodies shows endogenous localization of Fmn2 (grayscale in A and cyan in merge), actin (grayscale in A׳ and green in merge) and microtubule (grayscale A’’ and magenta in merge) and overlayed image of all three in (A’’’). Fmn2 decorates the entire filopodia and Fmn2-actin enriched filopodia are frequently occupied by microtubules as shown by arrowheads; Scale bar-5 μm. (B-B’’’) and (C-C’’’) representative STED micrographs of actin and microtubule in Ctl-MO and Fmn2-MO-treated growth cones, respectively. The alignment of microtubule (grayscale in B and C; magenta in merge) with actin (grayscale in B’ and C’; green in merge) occurs in regions peripheral from the center of the growth cone. An inset in (B’’’) shows alignment of microtubule and actin in lamellipodial region and filopodia in control and lack of such alignment in Fmn2-MO growth cone in (C’’’); Scale bar-5μm, inset-1 μm. (D) Quantification of filopodia number per growth cone (Ctl-MO, n=25; Fmn2-MO, n=28) and (E) growth cone filopodial lengths (Ctl-MO, n=168 and Fmn2-MO, n=118). (F) Percentage of filopodia per growth cone invaded by MT in Ctl-MO (n=24) compared to Fmn2-MO (n=27). (G) Number of MT ends per growth cone quantified in Ctl-MO (n=25) and Fmn2-MO (n=28) show that numbers are comparable even though the proportion of MT invading filopodia per growth cone is reduced in Fmn2-MO compared to control in (H) (Ctl-MO, n=24; Fmn2-MO, n=27). (I) The percentage of MT aligned to actin per growth cone in the peripheral zone (lamellipodial and filopodial regions) (Ctl-MO, n=24; Fmn2-MO, n=27) and the (J) lengths over which the actin and MT are co-aligned (Ctl-MO, n=161; Fmn2-MO, n=78) was much higher in control than Fmn2 depleted growth cones; ***P<0.001, **P<0.01, ns P>0.05; Mann-Whitney test.

Association and co-alignment with F-actin bundles have been proposed to guide MTs from the central region of the growth cone into the peripheral filopodia (Sabry et al., 1991, Schaefer et al., 2002, Zhou et al., 2002). We, therefore, evaluated the co-alignment of MTs with actin bundles in the peripheral zone (lamellipodial region and filopodia) of the growth cones. Fmn2 was found to promote alignment of MTs with F-actin bundles. The proportion of MTs aligned to actin bundles was drastically reduced from 71.61 ±4.44% in control neurons to 34.88 ±4.27% upon knockdown of Fmn2 (Figure 1 I). Furthermore, the distance over which the two polymers co-aligned also decreased upon reduction of Fmn2 expression (Figure 1 J).

These results implicate Fmn2 as a novel regulator of MT organization in growth cones. Our observations suggest Fmn2 function in mediating the association of MTs to F-actin bundles, thereby facilitating MT exploration of the peripheral zone and entry into filopodia.

### Microtubule dynamics and stability in growth cones is regulated by Fmn2

To examine Fmn2-dependent regulation of MTs in growth cones, we evaluated MT dynamics by imaging EGFP-tagged MT plus-end-tracking protein (+TIP), EB3. The *plusTipTracker* algorithm (Applegate et al., 2011, Matov et al., 2010) was used to quantify EB3 dynamics (Figure 2 A,B; Movie S1; Movie S2). Although the average number of dynamic MT tips in growth cones were comparable (Figure S2 A), the mean growth speed of MTs in the Fmn2 KD growth cones (8.49 ± 0.59 µm/min) was higher compared to controls (6.29 ± 0.45 µm/min; Figure 2 C). Consistent with this observation, the growth length of the polymerizing MTs also increased from 0.87 ± 0.07 µm in Ctl-MO transfected neurons to 1.17 ± 0.33 µm in Fmn2-MO -treated growth cones (Figure 2 D). However, the MT growth lifetimes remained unchanged (Figure 2 E; 7.67 ± 0.59 s in control and 7.83 ± 0.39 s in Fmn2 knockdown cells). Fmn2 knockdown resulted in an increase in the overall MT dynamicity (collective growth speeds of all comet tracks over their collective lifetimes; 7.45 ± 0.44 µm/min and 5.67 ± 0.39 µm/min in Fmn2-MO and Ctl-MO, respectively; Figure 2 F).

**Figure 2:**
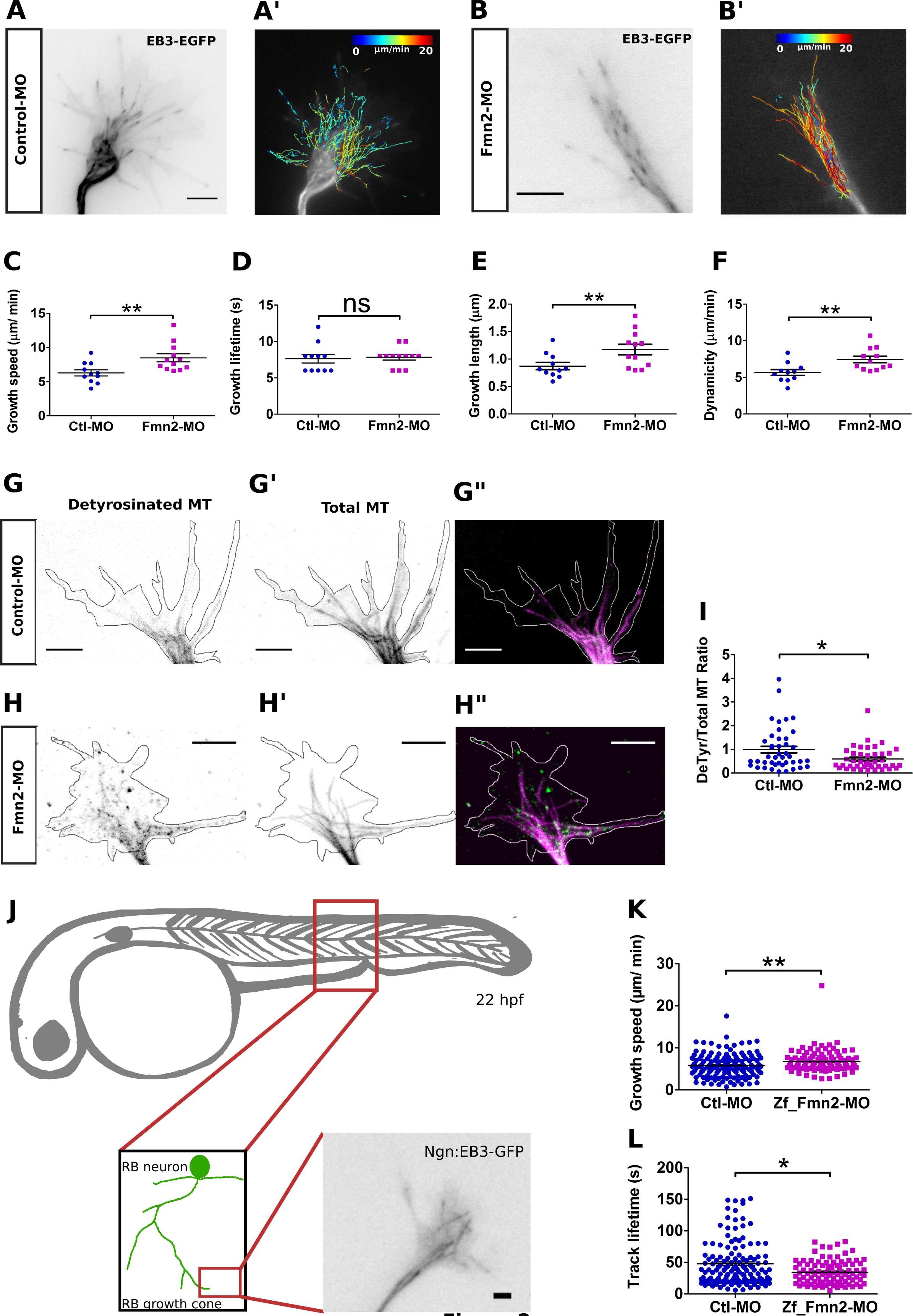
Knockdown of Fmn2 increases dynamicity and reduces stability of microtubules in the growth cone. (A and B) show representative growth cones labelled with the plus-tip marker GFP-EB3 in Ctl-MO and Fmn2-MO transfected growth cones, respectively. Time-lapse movies of EB3 were acquired at 2s interval for 100 frames to measure MT polymerization. The analysis was done using *plusTipTracker* software and the comet tracks with overlaid color-coded growth speeds are shown for Ctl-MO in (A’) and for Fmn2-MO in (B’); Scale bar-5μm. (C-F) show quantification of mean MT growth speed (C), mean MT growth lifetime (D), mean MT growth length (E), and overall dynamicity of MT polymerization in Ctl-MO (n=11; number of tracks = 2917) and Fmn2-MO (n=12; number of tracks = 2426) transfected growth cones; **P<0.01, ns P>0.05; Mann-Whitney test. (G -I) show representative images of detyrosinated (DeTyr) tubulin and total tubulin stained using specific antibodies for Ctl-MO (G and G’, respectively) and Fmn2-MO (H and H’, respectively). The intensity of detyrosinated tubulin was measured within the area occupied by the total tubulin signal and normalized to total tubulin intensity. This was quantified in (I) (n= 43 for Ctl-MO and n=42 for Fmn2-MO); Scale bar-5μm; *P<0.05; Mann-Whitney test. (J) Schematic representation of zebrafish larva at 22 hours post-fertilization (hpf) injected with Ngn1: EB3-GFP. EB3-GFP is expressed in the Rohon-Beard (RB) neurons (inset). The second inset shows a peripheral growth cone of a RB neuron displaying EB3 comets *in-vivo*. EB3 growth speeds (K) increase while the track lifetimes (L) decrease upon knockdown of zebrafish Fmn2 by specific morpholinos (Zf_Fmn2-MO). EB3 comets were analyzed using kymographs. (Ctl-MO, n=134 comet tracks from from 13 growth cones; Fmn2-MO, n=95 tracks from 12 growth cones); **P<0.01, *P<0.05; Mann-Whitney test.

Since the analysis of EB3 dynamics upon Fmn2 knockdown suggested an overall decrease in the stability of the MT cytoskeleton, we tested whether this observation was consistent with changes in the levels of detyrosinated tubulin – a tubulin post-translational modification associated with long lived, stable MTs. A mask of the area occupied by total tubulin immunofluorescence was used to limit the boundaries of the MT network and the fluorescence intensity of detyrosinated tubulin within this mask was quantified. Indeed, Fmn2 depletion reduced the detyrosinated α-tubulin/total MT ratio in growth cones (Figure 2 G-I).

Collectively, these data suggest that Fmn2 is an important regulator of MT dynamics in growth cones and promotes the stability of the MT network.

### Fmn2 regulates microtubule dynamics in growth cones *in vivo*

To evaluate the role of Fmn2 in growth cone MT dynamics *in vivo*, we expressed fluorescently tagged EB3 in the Rohon-Beard (RB) neurons of larval zebrafish.

Growth cones of the peripheral axons of the RB neurons expressing the reporter protein were evaluated in 22 hpf zebrafish embryos (Figure 2 J; Movie S3; Movie S4). A splice blocking morpholino against zebrafish Fmn2 (Zf_Fmn2-MO; see Materials and Methods for characterization; Figure S2 D) was used to deplete functional Fmn2. Compared to non-specific morpholino (Ctl-MO) injected animals, injection of Zf_Fmn2-MO resulted in increased growth speeds (Figure 2 K) and growth lifetimes (Figure 2 L) of EB3 comets in RB neuron growth cones.

These data indicate that depletion of Fmn2 *in vivo* alters MT dynamics in growth cones and is suggestive of a conserved MT regulatory function across phyla and neuronal subtypes.

### Microtubule stability in filopodia requires Fmn2

Exploratory MTs are guided into filopodia by co-alignment with F-actin bundles (Schaefer et al., 2002, Slater et al., 2019). As both the proportion of filopodia invaded by MT and the number of MTs entering filopodia were reduced upon Fmn2 knockdown (Figure 1 F,H), we evaluated the EB3 dynamics specifically within filopodia.

Growing dynamic MTs showed a characteristic reduction in velocity as they entered a filopodia from the central region of the growth cone, presumably due to retardation of growth speed resulting from direct interactions with filopodial actin bundles (Figure 3 A). However, in growth cones depleted of Fmn2, no change in the velocity was observed as MTs polymerised from the central region to enter filopodia (Figure 3 B). This observation suggests that Fmn2 may mediate the association of the invading MT with the filopodial F-actin bundle. Given the localization of Fmn2 along the entire length of the filopodial actin bundle, it is ideally positioned to guide exploratory MTs into filopodia. Consistent with these observations, in Fmn2 knockdown neurons, the growth speed of MTs inside filopodia was found to be higher (9.72 ± 0.52 µm/min) compared to control neurons (7.31 ± 0.29 µm/min; Figure 3 C). However, the lifetime of the EB3 comets within filopodia was reduced by 50% upon Fmn2 depletion (Figure 3 D; 86.40 ± 7.58 s in Ctl-MO and 40.91 ± 5.35 s in Fmn2-MO treated filopodia). The incursion distance of EB3 comets inside filopodia were unaltered between control and Fmn2-MO -treated neurons (Figure S3).

**Figure 3:**
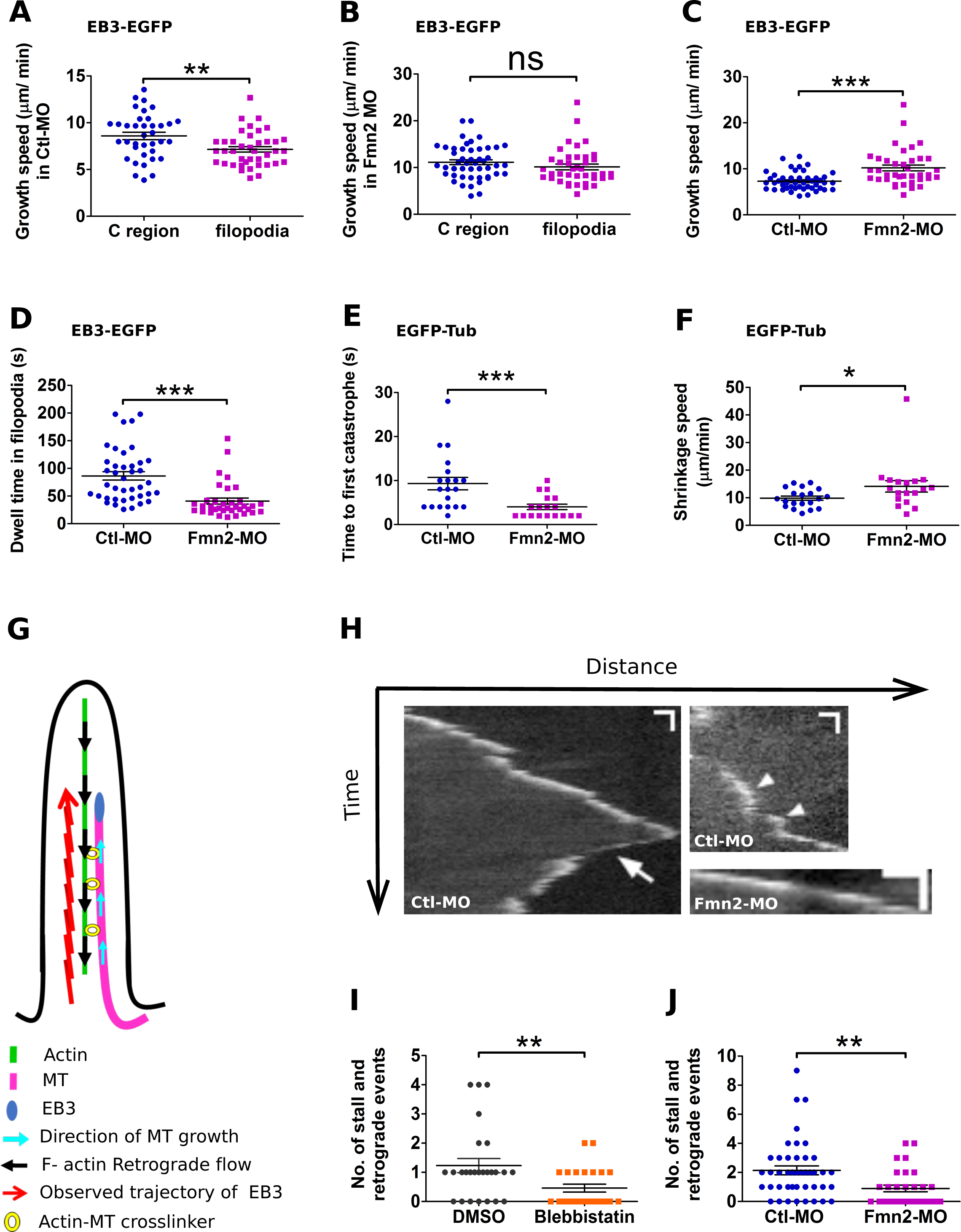
Fmn2 promotes MT stability in growth cone filopodia. (A and B) Growth speeds of EB3 comets in the transition zone and filopodial region of Ctl_MO (T zone, n=37 and filopodia, n=46) and Fmn2-MO (T zone, n=37 and filopodia, n=38) treated growth cones, respectively. (C and D) Comparison of growth speeds (Ctl-MO, n=46 and Fmn2-MO, n=38) and dwell time of EB3 (Ctl-MO, n=40 and Fmn2-MO, n=35) comets inside filopodia, respectively. (E and F) Quantification of time to first catastrophe after entering filopodia (Ctl-MO, n=20 and Fmn2-MO, n=18) and catastrophe speed (Ctl-MO, n=20 and Fmn2-MO, n=18), respectively, measured using EGFP-tubulin in Ctl-MO and Fmn2-MO filopodia. (G) schematic showing MT growth and observed trajectory of EB3 comets along actin bundles inside filopodia. The trajectory of EB3 comets appear jagged with retrograde and stall events. This is due to the coupling of the growing MT filament with F-actin, which itself is experiencing centripetally directed retrograde flow. (H) Representative kymographs of EB3 showing retrograde (arrow) and stall (arrowhead) events in Ctl-MO and Fmn2-MO treated filopodia. Fmn2 knock down decoupled the growing MT from the filopodial F-actin and resulted in smoother trajectories, as seen in Fmn2-MO kymograph. (I) Quantification of retrograde and pause events for blebbistatin treatment (n=26, DMSO and n=24, blebbistatin) used to reduce F-actin retrograde flow. (J) Reduction in stall and retrograde events in EB3 trajectories upon depletion of Fmn2 (Ctl-MO, n=43; Fmn2-MO, n=30 neurons); Scale bars-10μm, 10s; ***P<0.001, **P<0.01, ns P>0.05; Mann-Whitney test.

The reduction in the dwell time of EB3 comets in filopodia is suggestive of loss of MT stability. We probed this further by measuring MT catastrophe events directly using fluorescently-tagged α-tubulin. This analysis revealed that MT catastrophe was more frequent upon Fmn2 knockdown as the latency for the first catastrophe event following incursion into a filopodia was much shorter (Figure 3 E; 4.00 ± 0.61s for Fmn2-MO and 9.30 ± 1.42s for Control-MO). The propensity for catastrophe possibly accounts for the short-lived dwell times of EB3 comets in filopodia. Further, the velocity of retraction/shrinkage during catastrophe of MTs within filopodia in Fmn2-MO transfected neurons (14.13 ± 2.08 µm/min) was significantly higher than the Ctl-MO treated filopodia (9.87 ± 0.76 µm/min) (Figure 3 F). The latter observation is consistent with Fmn2-dependent association between MT and filopodial F-actin that prevents rapid retraction of a collapsing MT filament.

Filopodial F-actin bundles undergo retrograde flow and the invading MT, owing to its association with these actin bundles, also experiences this retrograde movement (Schaefer et al., 2002). This behaviour can be captured in kymographs of EB3 comets inside filopodia as sporadic rearward movement (retrograde events) and apparent cessation of forward progress (stall) despite ongoing MT polymerization as evident by the persistence of the EB3 signal at the tip of the invading MT (Figure 3 H) (Liu et al., 2010). As expected, the reduction of F-actin retrograde flow by the myosin-II inhibitor blebbistatin made the EB3 trajectories smoother and decreased the EB3 retrograde and stalls (Figure 3 I). Significantly, knockdown of Fmn2 resulted in the reduction of retrograde and stall events and much smoother MT growth trajectories inside filopodia (Figure 3 J) indicating the involvement of Fmn2 in mediating the association of MTs with filopodial F-actin.

Taken together, these data indicate that Fmn2 not only mediates the capture and guidance of exploratory MTs into filopodia, but also regulates the stability of the invasive MT within the filopodia. Thus, Fmn2, directly or indirectly, crosslinks the polymerising MTs to the filopodial F-actin bundles.

### Fmn2 directly binds and bundles F-actin and Microtubules

*In vitro* studies on the fly formin, Capuccinno (Capu), indicate that the Capu C-terminal FSI domain mediates charge-based interaction with both F-actin and MTs (Roth-Johnson et al., 2014). Alignment of Capu and chick Fmn2 FSI domains revealed a high degree of homology including the conservation of the basic amino acid residues mediating the charge-based interactions of Capu with F-actin and MTs (Figure S4 A). To directly probe the interactions of Fmn2 with actin and MTs, we purified recombinant proteins corresponding to the FH2FSI and FH2_ΔFSI regions of the chick Fmn2 protein (Figure 4 A,B). The actin binding activities of these proteins were tested using a conventional high-speed co-sedimentation assay. FH2FSI and FH2_ΔFSI bound to actin filaments with apparent equilibrium constants (Kd) of 5.0 ± 0.2 µM and 12.5 ± 0.1 µM, respectively (Figure 4 C, S4 B,C). These findings demonstrate higher affinity of FH2FSI for F-actin than FH2_ΔFSI.

**Figure 4:**
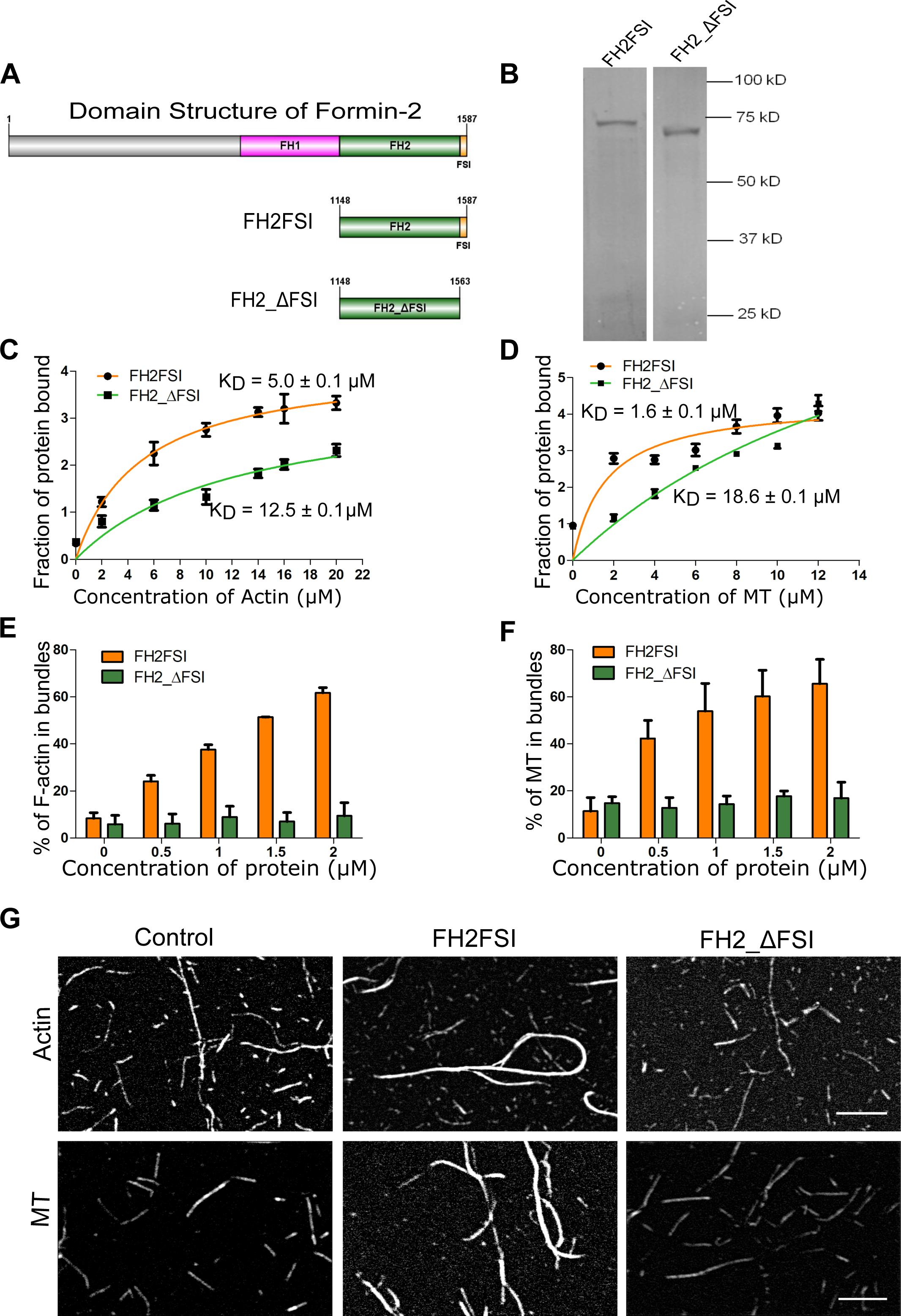
Fmn2 can interact with and bundle both actin and MT. (A) Schematic representation of the domain structure chick Fmn2 and the different constructs used in this study. (B) Representative gel showing the expression of purified FH2FSI and FH2_ΔFSI constructs used for the study. (C) Binding of FH2FSI and FH2_ΔFSI to F-actin. Increasing concentrations of F-actin was incubated with 2 µM of each construct. The percentage of bound proteins from each constructs was plotted against F-actin concentrations and fitted with a hyperbolic function. The Kd for FH2FSI and FH2_ΔFSI are 5.0 ± 0.1 µM and 12.5 ± 0.1 µM, respectively (mean ± SD from three independent experiments). (d) FH2FSI and FH2_ΔFSI binding affinity for MT. Different concentrations of MT was incubated with 2 µM of each construct. The amount of protein from each construct bound to microtubule was fitted to hyperbolic function for three independent reactions. The Kd for FH2FSI and FH2_ΔFSI are 1.6 ± 0.1 µM and 18.6 ± 0.1 µM, respectively (mean ± SD from four independent experiments). (E) Quantification of the actin bundles in presence of FH2FSI and FH2_ΔFSI from low speed sedimentation assays for F-actin bundling. The data represents the mean ± SEM from three independent experiments. (F) Quantification of microtubule bundling by FH2FSI and FH2_ΔFSI from low speed sedimentation assays for MT bundling. The data represents the mean ± SEM from three independent experiments. (G) Representative TIRF microscopy images of the F-actin (upper panel) and MT (lower panel) bundling reactions after incubation of 1 μM FH2FSI and 1 μM FH2_ΔFSI with sparsely labelled F-actin or MT for 15 mins. Scale bar, 5 μm.

We further investigated the MT binding ability of FH2FSI and FH2_ΔFSI in a high-speed co-sedimentation assay. Our data revealed that Fmn2 FH2FSI has significantly higher affinity for taxol-stabilized MTs than FH2_ΔFSI with Kd of 1.6 ± 0.1 µM and 18.6 ± 0.1 µM, respectively (Figure 4 D; S4 D,E). Thus, Fmn2 is capable of binding both F-actin and MTs directly and that the FSI domain is a key component of these associations.

Apart from binding F-actin, FH2FSI was also capable of bundling F-actin filaments as demonstrated by low speed co-sedimentation assays and total internal reflection fluorescence (TIRF) microscopy of labelled F-actin filaments (Figure 4 E,G, S4 F). In contrast, FH2_ΔFSI was incapable in bundling actin filaments (Figure 4 E, G, S4 G).

Low speed co-sedimentation with taxol-stabilized MTs revealed that FH2FSI, but not FH2_ΔFSI, could also bundle MTs in a concentration-dependent manner (Figure 4 F; S4 H,I). This observation was confirmed using TIRF imaging of labelled MTs in the presence of recombinant proteins (Figure 4 G).

Further, we tested the ability of Fmn2 to protect MTs from cold shock-induced depolymerization. FH2FSI robustly stabilized MTs from implicating a role in protecting MTs from depolymerisation (Figure S4 J).

These *in vitro* studies suggest that Fmn2 is a direct interactor of both F-actin and MTs and the FSI domain is a major contributor to these interactions. Moreover, the FSI domain was found to be necessary for efficient bundling F-actin and MT filaments.

### F-actin and microtubule filaments are cross-linked by Fmn2

The ability of Fmn2 to bind both F-actin and MTs led us to directly test the ability of Fmn2 to crosslink these two filament types. We visualised fluorescently-labelled pre-polymerised phalloidin stabilized F-actin and taxol-stabilized MT filaments with and without FH2FSI or FH2ΔFSI. In controls lacking Fmn2 fragments, randomly oriented single filaments of F-actin and MT were observed (Figure 5 A). However, in the presence of FH2FSI, F-actin and MT filaments co-aligned to form thick hybrid bundles (Figure 5 A). On the other hand, the deletion of the FSI domain (FH2ΔFSI) resulted in lack of co-alignment and filaments were randomly distributed as seen in no protein controls (Figure 5 A).

**Figure 5:**
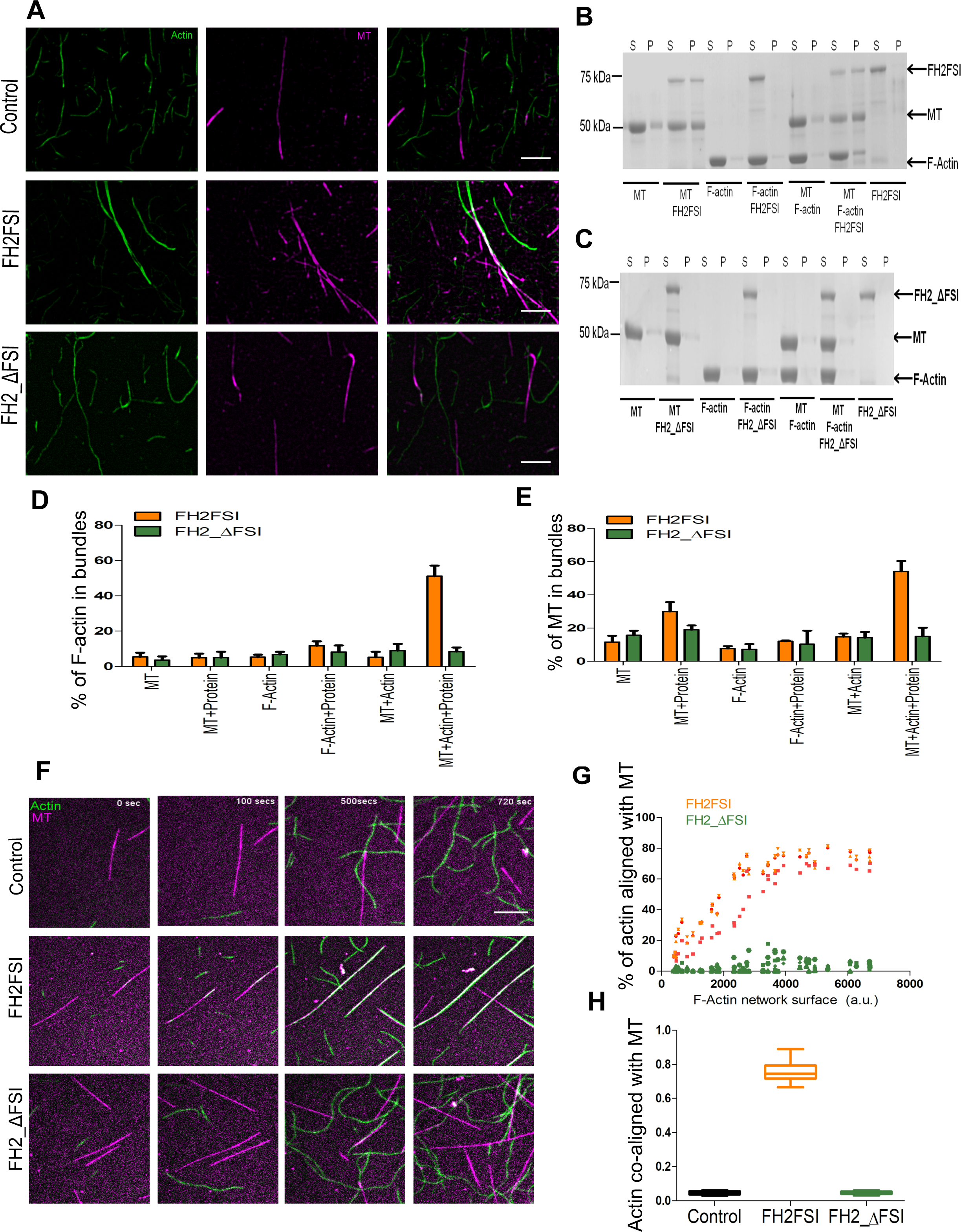
The FSI domain of Fmn2 is required for efficient crosslinking of the F-actin and microtubules. (A) Visualization of the phalloidin-stabilized actin and taxol-stabilized microtubule cross-linking in absence or purified protein (upper panel; control) or in the presence of FH2FSI (middle panel) or FH2_ΔFSI (lower panel). The reaction mixture was incubated with the FH2FSI and FH2_ΔFSI for 30 mins before centrifugation followed by visualisation. Scale bar, 5 μm. (B) Representative SDS-PAGE showing cross-linking of phalloidin-stabilized actin and taxol-stabilized microtubule polymers by FH2FSI after incubation for 30 mins and centrifugation at 2,000g. (C) Representative SDS-PAGE showing lack of cross-linking of the actin and microtubule complexes in the presence of FH2_ΔFSI after incubation for 30 mins and centrifugation at 2,000g. (D) Quantification of the cross-linked phalloidin-stabilized actin in presence of FH2FSI or FH2_ΔFSI from the co-sedimentation assays. The data represents the mean ± SEM from four independent experiments. (E) Quantification of the cross-linked taxol-stabilized microtubules in presence of FH2FSI or FH2_ΔFSI from the co-sedimentation assays. The data represents the mean ± SEM from four independent experiments. (F) Representative frames from time lapse TIRF imaging of co-polymerizing microtubules and F-actin. In the control (upper panel) 25 μM tubulin dimers (magenta) was co-polymerized with 1 μM actin monomers (green). The filaments were distributed randomly without any association. Addition of FH2FSI (middle panel) resulted in the co-assembling the MT and F-actin filaments to co-align and form thick hybrid bundles. However, addition of FH2_ΔFSI (lower panel) did not align and cross-bridge MT and F-actin and each of the individual filaments remained un-associated with each other and were distributed randomly. Scale bar, 10 μm. (G) Percentage of F-actin co-aligned with microtubule as a function of the F-actin network surface in the presence of FH2FSI or FH2_ΔFSI (four individual curves for each protein). a.u., represents arbitary units. (H) Box plot analysis of the F-actin surface co-aligned with microtubules in presence FH2FSI and FH2_ΔFSI (the graph represents the co-alignment at the end of 720 secs of 20 individual data sets in each condition).

F-actin-MT crosslinking ability of Fmn2 was further assessed using sedimentation assays. FH2FSI or FH2ΔFSI was incubated with pre-polymerised phalloidin stabilized F-actin and/or taxol-stabilized MTs and centrifuged at very low speeds. The centrifugation conditions were optimised such that incubation of FH2FSI with MT or F-actin alone resulted in partitioning only a small amount of the polymers in the pellet fraction. However, incubation of FH2FSI with F-actin and MTs simultaneously resulted in a much larger amount of both the cytoskeleton polymers partitioning into the pellet fraction (Figure 5 B,D-E). These data indicate that Fmn2 interacts with both F-actin and MTs filaments simultaneously and results in the formation of a larger hybrid complex. Incubation with FH2ΔFSI failed to show similar crosslinking between MTs and F-actin (Figure 5 C-E).

To directly and dynamically visualise the formation of hybrid complexes and co-alignment between F-actin and MTs, we monitored the co-polymerisation of the two cytoskeletal polymers using TIRF microscopy. MT and actin filaments were allowed to self-polymerize and elongate using tubulin dimers and actin monomers (a fraction of the tubulin dimers and actin monomers were fluorescently labelled). In controls lacking Fmn2 protein, both F-actin and MT filaments were found to elongate but rarely contacted or associated with each other (Movie S5). Strikingly, the presence of FH2FSI induced progressive co-alignment of the extending F-actin and MT filaments (Figure 5 F-H; Movie S6). The association of the relatively more flexible F-actin filaments with the stiffer MTs resulted in progressive straightening of the F-actin filaments. In contrast, FH2ΔFSI failed to co-align the two filaments underscoring the centrality of the FSI domain in simultaneous binding to F-actin and MTs by Fmn2 dimers to promote crosslinking (Figure 5 F-H; Movie S7).

### Actin-microtubule crosslinking by Fmn2 is necessary for microtubule stability in filopodia

As our *in vitro* studies demonstrated the requirement of the FSI domain in actin-MT crosslinking by Fmn2, we used the presence or absence of this domain in rescue experiments to directly probe Fmn2 function in growth cone filopodia. The cDNA for the mouse orthologue of Fmn2 (mFmn2) lacks the binding sequence for anti-chick Fmn2 morpholinos and is thus resistant to knockdown by this reagent. EB3 dynamics within filopodia was evaluated following co-transfection with Fmn2-MO/Ctl-MO and morpholino-resistant mFmn2 constructs. Expression of the full-length mouse Fmn2 cDNA (mFmn2_FL), in the background of endogenous chick Fmn2 knockdown, was able to rescue the defects observed in EB3 comet speed (Figure 7 A), dwell time (Figure 7 B) and the EB3 retrograde/stall events (Figure 7 C) inside filopodia. However, mouse Fmn2 lacking the FSI domain (mFmn2_ΔFSI) failed to rescue all of these parameters (Figure 7 A-C).

**Figure 6:**
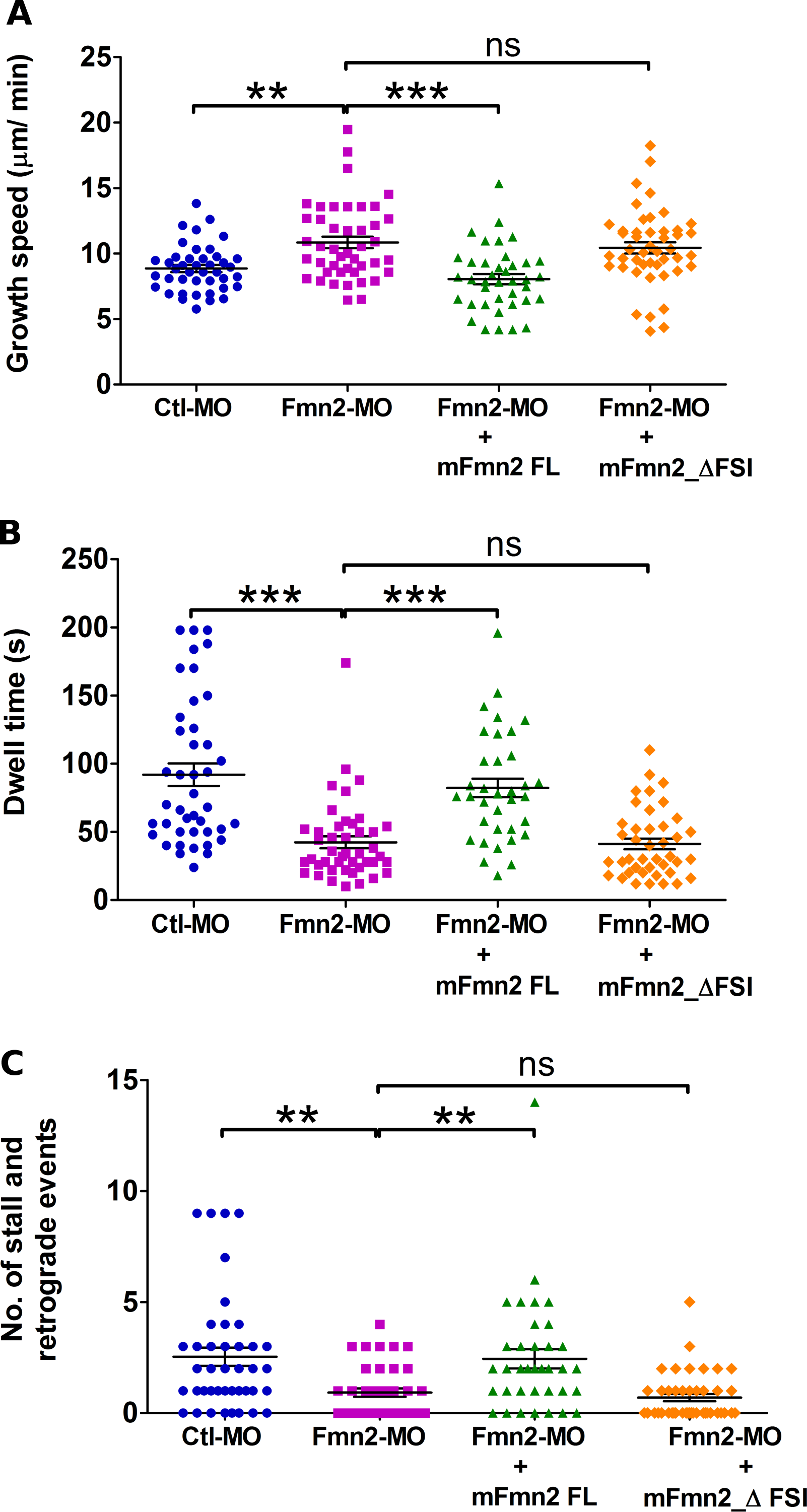
The FSI domain of Fmn2 is required for coupling actin and MT in filopodia. (A) Growth speed, (B) dwell time and (C) retrograde and pause events of EB3 comets inside filopodia were rescued by morpholino-resistant full-length mouse Fmn2 but not when the FSI domain was lacking. (Ctl-MO, n=42; Fmn2-MO, n=44; mFmn2-FL rescue, n = 35; mFmn2_Δ_FSI,n= 41; for graph A. Ctl-MO, n=42; Fmn2-MO, n=44; mFmn2-FL rescue, n = 39; mFmn2_Δ_FSI, n= 46; for graph B.; Ctl-MO, n=41; Fmn2-MO, n=39; mFmn2-FL rescue, n = 36; mFmn2_Δ_FSI, n= 43 for graph C); ns > P .0.05; **P<0.01; ***P<0.001; Krusal-Wallis test followed by Dunn’s multiple comparison test.

**Figure 7:**
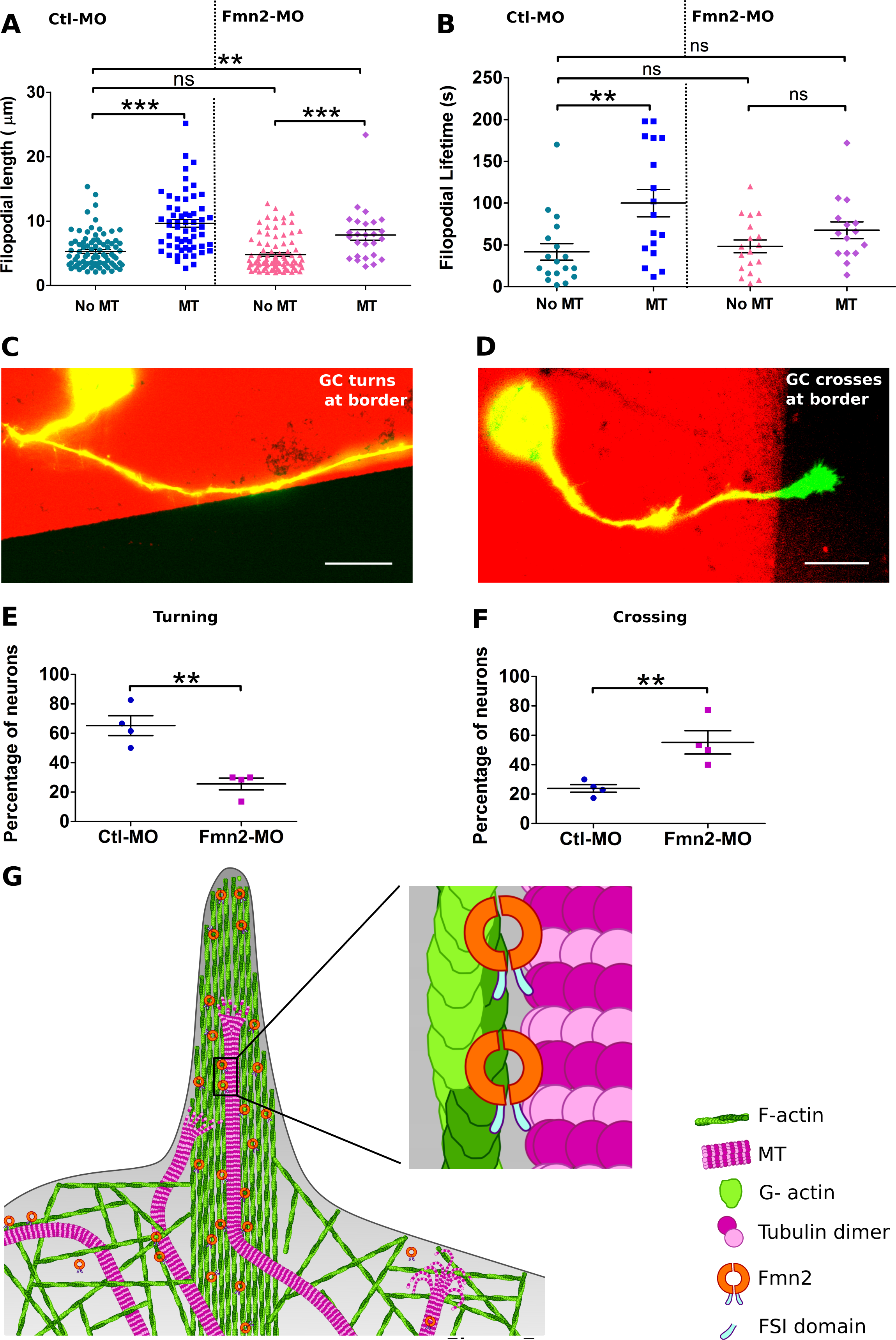
Fmn2 is required for filopodial stability and growth cone turning *in-vitro* and MT stability *in-vivo*. (A) Comparison of filopodial lengths with and without MT inside from Ctl-MO and Fmn2-MO treated growth cones. Immunofluorescence was used to detect MTs while labelled phalloidin marked F-actin (Ctl-MO + No MT, n=92; Ctl-MO + MT, n= 104; Fmn2-MO + No MT, n=83; Fmn2-MO + MT, n=21). (B) Comparison of filopodial lifetimes with or without EB3 entry during the time period of imaging - 100 frames with 2 sec interval (Ctl-MO + No MT, n=18; Ctl-MO + MT, n= 17; Fmn2-MO + No MT, n=18; Fmn2-MO + MT, n=15). ns P >0.05; *P<0.05, **P<0.01; ***P<0.001; Krusal-Wallis test followed by Dunn’s multiple comparison test. (C) Representative image of a control neuron co-transfected with Ctl-MO and soluble GFP showing a growth cone turning at the border between Laminin + Fibronectin (L + FN) and poly-l-lysine (PLL) to continue to stay on the preferred L + FN region. (D) representative image of Fmn2-MO and GFP co-transfected growth cone failing to turn and crossing over from the L + FN side into the PLL region. Quantification of turning (E) and crossing (F) events of growth cones from the L + FN region encountering a border with PLL (Ctl-MO=68; Fmn2-MO=78; data compiled from 4 independent biological replicates). Scale bar-20μm, **P<0.01; Unpaired t test. (G) Schematic representation of a growth cone filopodium with parallel actin bundles showing microtubule incursion into the filopodium. Fmn2 dimers on filopodial F-actin bundles cross-bridge exploratory microtubules resulting in co-alignment and targeting into filopodia. Inset suggests a model where one monomer of the Fmn2 dimer binds F-actin and the other binds to the microtubule lattice. Fmn2-dependent crosslinking of MT to F-actin in filopodia increases the stability of the invading MT. The filopodial lifetimes and stability are increased due to the stable MT providing a providing a track for cargo delivery and increasing the mechanical resilience to the filopodium. It is proposed that biased stabilization of filopodia, in response to chemo/hapto-tactic cues result in growth cone turning.

These rescue experiments demonstrate that, in agreement with our *in vitro* studies, Fmn2 regulates MT dynamics in filopodia by crosslinking MTs to the filopodial F-actin via the FSI domain.

### Microtubule entry stabilizes filopodia

MT entry and capture within filopodia is known to stabilize filopodia and allow its extension (Barzik et al., 2014, Hu et al., 2012, McNeely et al., 2017). We tested the association between the presence of MTs within growth cone filopodia and filopodial lengths and lifetimes.

We probed association between the presence MT within filopodia and filopodial dynamics. Comparison of filopodial populations with and without MTs revealed a strong association between the presence of MT and filopodial stability. In Ctl-MO treated growth cones, immunofluorescence analysis using labelled phalloidin and antibodies against α-tubulin revealed that filopodia with detectable MTs had longer lengths (9.64±0.59 µm) compared to those lacking MTs (5.30±0.29 µm; Figure 7 A). Live imaging studies tracking EB3 further concluded that filopodia associated with MTs had longer lifetimes (100.0±16.33 s) than those without MT (41.67± 9.85 s; Figure 7 B). Importantly, the same trend is seen in Fmn2 depleted neurons. The filopodia not containing MT are shorter (4.81±0.28 µm) and also short-lived (48.22 ± 7.71 s) compared to filopodia with MTs (7.84 ±0.81 µm and 67.60 ± 10.04 s) (Figure 7 A,B).

These results suggest that the reduced stability of MTs in filopodia (Figure 3 D-F), and the consequent reduction in the number of filopodia having MTs at any given time (Figure 1 F) is a major contributor to the reduced filopodial lengths and lifetimes observed in Fmn2 depleted growth cones (Sahasrabudhe et al., 2016).

### Fmn2 is required for growth cone turning at substrate borders

The deficits in MT entry into filopodia and the consequent reduction in filopodia stability is likely to influence chemotactic response of Fmn2 depleted growth cones (Williamson et al., 1996, Zheng et al., 1996). We tested this proposition using a haptotactic border turning assay (Jean et al., 2012, Liu et al., 2010, Turney & Bridgman, 2005).

Neurons were cultured on alternating stripes of fibronectin + laminin and poly-l-lysine (PLL) generated by microcontact printing and evaluated for growth cone turning behaviour at substrate borders (Figure 7 C, D). Although both surfaces were independently permissive for neuronal growth, growth cones of neurons plated on laminin + fibronectin preferred to turn and continue to reside on the same substrate upon encountering the border with PLL. In Ctl-MO transfected neurons, ∼73% of growth cones turned upon encountering the border (Figure 7 E-F). However, upon depletion of Fmn2, only ∼32% turned while the majority crossed the border to grow into PLL (Figure 7 E-F).

These observations indicate aberrant chemotactic behaviour of Fmn2 knockdown growth cones possibly originating from deficits in MT stabilization in filopodia resulting in the inability to execute reliable growth cone turning at substrate borders.

## DISCUSSION

Coordination and co-regulation of the actin and MT cytoskeletons are essential for key cellular processes, including motility and pathfinding of neuronal growth cones. However, our understanding of molecular activities mediating this crosstalk is limited. We identify Fmn2 as a novel mediator of actin-MT coordination in neuronal growth cones. Using a combination of *in vitro* biochemistry and cell biological analysis of spinal growth cones, we show that Fmn2 can directly interact and crossbridge F-actin and MTs and this function is critical for growth cone dynamics and haptotaxis.

In recent years, several lines of evidence have accumulated implicating Fmn2 function in neurodevelopment and neurodegeneration (Agis-Balboa et al., 2017, Almuqbil et al., 2013, Law et al., 2014, Marco et al., 2018). However, little is known about the mechanistic underpinnings of Fmn2 function in the nervous system. We have previously shown that Fmn2-depleted spinal commissural axons fail to cross the floor plate and show aberrant trajectories post crossing (Sahasrabudhe et al., 2016). Axonal pathfinding defects and deficits in the dynamics of chemosensory filopodia described in this earlier study prompted us to evaluate Fmn2 function in coordinating MT-actin dynamics in the growth cone.

We found that Fmn2 was necessary to stabilize MT dynamics in the growth cone. Depletion of Fmn2 resulted in increased MT dynamicity and polymerisation accompanied by a reduction in the population of stable, long lived detyrosinated MTs.

STED nanoscopy revealed that co-alignment of F-actin bundles and MTs was reduced in Fmn2 knockdown growth cones. This observation implicates Fmn2 in regulating interpolymer association between actin and MT. Thus, the loss of association with F-actin may result in the observed changes in MT dynamics in the growth cone. Additionally, the ability of Fmn2 to bundle MT filaments and protect MTs from cold-shock-induced depolymerisation are also likely to directly influence MT dynamics.

As F-actin bundles are known to guide MTs into filopodia via direct or indirect interactions (Schaefer et al., 2002, Zhou et al., 2002), we conducted a detailed analysis of MT dynamics inside filopodia. Our data reveal two phenomena. First, Fmn2 along filopodial actin bundles mediate dynamic associations with exploratory MTs and guide them into filopodia (Figure 7). Similar capture and redirection of MTs by F-actin bundles have previously been demonstrated *in vitro* (Preciado Lopez et al., 2014). Second, in Fmn2 knockdown cells, despite a faster growth rate within the filopodia (possibly reflecting reduced dynamic interactions with F-actin), the invading MTs are unstable. Both the reduced dwell time of EB3 comets and increased predisposition to catastrophe as seen in the GFP-tubulin studies indicate reduction in stability.

We directly tested the F-actin-MT cross bridging activity of Fmn2 using *in vitro* biochemistry and reconstitution strategies with purified recombinant Fmn2 fragments. These studies revealed that the Fmn2 dimer is a potent actin-MT crosslinker with its C-terminal FSI domain central to this function. The conserved FSI domain of the *Drosophila* ortholog of Fmn2, Capu, has been previously shown to interact with both F-actin and MTs via non-specific charge-based interactions (Roth-Johnson et al., 2014) and Capu has been implicated in actin-MT crosslinking in fly oocytes (Dahlgaard et al., 2007, Rosales-Nieves et al., 2006). In neurons, expression of full-length Fmn2 but not FSI domain deleted constructs rescued the MT phenotypes associated with Fmn2 depletion underscoring the importance of this domain in microfilament-MT crosslinking.

Taken together, our data posits crosslinking activity as the central mechanism by which Fmn2 regulates MT dynamics in filopodia. It is possible that there are additional mechanism(s) involved. For example, Fmn2 depletion may result in the sparsening of filopodial F-actin contributing to the observed phenotypes. A contrary argument is that a reduction in physical hindrance to MT advance should have reduced predisposition to MT catastrophe and not increased it. Further, similar incursion depths of MTs in control and Fmn2 depleted filopodia also suggests mechanical hindrance to MT extension is unlikely to have been altered. However, our *in vitro* studies provide compelling support for the cross-bridging mechanism to play a significant role.

The FSI domain is the common interface between both F-actin and MT binding and it is also required for efficient actin nucleation/elongation by the FH2 domain (Vizcarra et al., 2014). Consequently, it is not possible to examine the effect of Fmn2 on actin and MT dynamics independently. *In vitro* studies suggest that MTs can inhibit actin nucleation by Capu but does not influence activity when Capu is bound to the growing barbed ends of F-actin filaments (Roth-Johnson et al., 2014). We reason that the FSI domain is involved in supporting nucleation/elongation F-actin when bound to the F-actin barbed end. However, it is also capable of F-actin filament side binding via non-specific charge interactions. The latter activity results in attenuated actin nucleation activity. When an invading MT reaches Fmn2-decorated filopodial F-actin bundles, one FSI domain of the Fmn2 dimer attaches to the polymerizing MT while the other FSI domain continues to interact with F-actin. This coupling guides the MT into the filopodia and stabilizes it from catastrophe (Figure 7 G). Growth cone turning involves selective stabilization of the chemosensory filopodia towards the direction of the turning (Zheng et al., 1996) which is facilitated by the increased capture of MTs within filopodia (Buck & Zheng, 2002, Mack et al., 2000, Sabry et al., 1991, Williamson et al., 1996). Stable MT within filopodia ensures targeted cargo delivery and also enhances the mechanical resilience of the filopodia. Consistent with this model, we find that MT entry is associated with longer filopodial lifetimes and lengths (Barzik et al., 2014, McNeely et al., 2017). As Fmn2 mediates the capture, incursion and lifetimes of MT inside filopodia, we find that depletion of Fmn2 results in deficits in growth cone turning at substrate boundaries.

As Fmn2 is associated with intellectual disability, learning and memory deficits and is localized to dendritic spines (Law et al., 2014), it will be of interest to examine the role of Fmn2 mediated actin-MT crosstalk in the context of structural plasticity of dendritic spines (Koganezawa et al., 2017, Merriam et al., 2013). Similarly, the enhanced dynamicity of MTs may have implications on axonal re-growth following injury (Blanquie & Bradke, 2018, Tang & Chisholm, 2016).

In oocytes, a Fmn2-dependent F-actin “cage” is assembled around the MT spindle and is essential for spindle positioning and chromosome segregation (Azoury et al., 2008, Mogessie & Schuh, 2017). It is tempting to speculate that actin-MT crosstalk facilitated by Fmn2 may regulate the development of the F-actin “cage” and also explain the reduction in kinetochore fibres in Fmn2-/- oocytes (Mogessie & Schuh, 2017).

In summary, this study identifies Fmn2 as a novel mediator of actin-MT crosstalk and provides a mechanistic framework for uncovering neurodevelopmental functions of Fmn2. Our study underscores the criticality of precise coordination between the actin and MT cytoskeletons in highly precise and adaptive processes like chemotactic motility of cells.

## MATERIALS AND METHODS (redo references)

### Plasmid constructs

The details of the source and cloning of plasmids used in this study are listed in Table S1. Neurons were co-transfected with pCAG-GFP or pCAG-mCherry plasmids along with morpholinos to select for transfected growth cones. pCAG-EB3-GFP, pCAG-EB3-mCherry or pCS2-GFP-tubulin was used to visualize MT dynamics.

Ngn1: EB3-GFP and Ngn1: TagRFP-CAAX was used to label polymerizing MTs and label the membrane of zebrafish RB neurons. Rescue experiments were undertaken with pCAG-mFmn2-GFP and pCAG-mFmn2-ΔFSI-GFP. For expression of recombinant protein fragments in bacteria, chick Fmn2 FH2FSI (1148^th^-1587^th^ aa) and FH2_ΔFSI (1148^th^-1563^rd^ aa) fragments were generated from full length chick Fmn2 (GenBank: KU711529.1) and cloned into pGEX-6P1 vector (gift from Dr. Thomas Pucadyil, IISER Pune).

### Primary neuronal cultures and transfection

Freshly fertilized, White Leghorn eggs were obtained from Venkateshwara Hatcheries Ltd., India. All protocols were followed in accordance with procedures approved by the Institutional Animal Ethics Committee, IISER Pune, India. HH stage 25-26 embryos were dissected in a laminar flow hood and spinal tissue between the fore and hind limbs was isolated. The tissue was dissociated using trypsin (0.25% trypsin + EDTA; Lonza) by incubation at 37 °C for 15 minutes. The dissociated tissue, then was electroporated with suitable plasmid (5-20µg) and/or morpholinos (100µM) in Optimem® (Gibco) media and plated in culturing media (L-15, 10% FBS and 1x Pen-Strep; all Gibco) and allowed to grow at 37 °C without CO₂ for 24 to 36 hours. Before plating, the glass coverslips were coated with 1 mg/ml poly-L-lysine (PLL; Sigma) and 20 µg/ml laminin (Sigma).

### Morpholino-based knockdown

To knockdown Fmn2 in chick neurons, anti-Fmn2 translation-blocking morpholinos (5’ CCATCTTGATTCCCCATGATTTTTC 3’) were used along with non-specific morpholinos as a negative control (5’ CCTCTTACCTCAGTTACAATTTATA 3’).

Previously, we have characterised this and other morpholino sequences and demonstrated effective and specific knockdown of Fmn2 protein expression in chick spinal commissural neurons (Sahasrabudhe et al., 2016). This was established by immunoblot and immunofluorescence analysis employing antibodies against endogenous chick Fmn2. In this study, we report greater than 75% knockdown of endogenous protein in growth cones of spinal neurons using immunofluorescence (Figure S1A). In these experiments, pCAG-GFP plasmid was co-tranfected along with morpholinos and incubated for 36 hours before immunostaining with anti-Fmn2 (custom made in the lab; Sahsrabudhe et al., 2016). Only GFP-expressing neurons were considered for analysis. Fmn2 fluorescence was normalized to growth cone area.

Specificity of this morpholino has been established by rescuing the knockdown phenotypes by co-expressing morpholino-resistant Fmn2 cDNA in primary neurons (this study and (Sahasrabudhe et al., 2016)) and *in vivo* (Sahasrabudhe et al., 2016).

### Immunofluorescence

Neurons were cultured for 24 hours before fixing. For STED and detyosinated / total tubulin ratio imaging, staining was done according to the procedure presented in (Biswas & Kalil, 2018). Briefly, the cultures were pre-extracted for 90 seconds with 0.4% glutaraldehyde +0.2% Triton X-100 in PHEM buffer (60mM PIPES, 25mM HEPES, 10mM EGTA, and 4mM MgSO4·7H20; all sourced from Sigma) and then fixed in 3% glutaraldehyde (in PHEM buffer) for 15 minutes in 37°C. After washes with 0.5% BSA in 1X PBS, permeabilization was performed with 0.5% Triton X-100 for 10 minutes at room temperature (RT). Glutaraldehyde was quenched with 5 mg/ml sodium borohydride for 5 minutes in 1x PBS. Blocking buffer of 10% BSA was used for 1 hour. Primary antibody incubation was done overnight at 4°C, followed by secondary antibody incubation for 1 hour at RT. The sample were then mounted in Mowiol-DABCO medium (2.5% 1,4-Diazabiocyclo-octane (DABCO), 10% Mowiol 4-88, 25% glycerol and 0.1M Tris-HCL, pH-8.5; all sourced from Sigma). For, triple antibody staining of Fmn2, MT and actin, the neurons were fixed with 4% paraformaldehyde and 0.05% glutaraldehyde in PHEM buffer for 20 minutes at RT. The neurons were permeabilised with 0.01% Triton X-100 for 10 minutes. Incubation with primary antibody was done in blocking buffer (3% BSA in 1X PHEM) for at 4°C overnight. Secondary antibody incubation was kept for 1 hour before mounting in 80% glycerol. All the washes were done in 1x PHEM buffer. The primary antibody for staining total MT (anti-α-tubulin, DM1a, sigma) and detyrosinated tubulin (EMD Millipore, AB3201) was at 1:3000 and 1:500 dilution, respectively. Anti-Fmn2 antibody was generated in house (Sahasrabudhe et al., 2016) and was used at 1:200 dilution. Secondary antibodies from Invitrogen, namely, Goat anti-mouse IgG Alexa Fluor 568 and Goat anti-mouse IgG Alexa Fluor 405 (for total MT) were used at 1:1000 dilution. While, Goat anti-rabbit IgG Alexa Fluor 488 (for detyrosinated MT and Fmn2; Invitrogen) was used at 1:500 dilution. F-actin was visualized with Alexa Fluor 633 phalloidin and Alexa Fluor 568 phalloidin (Invitrogen) at 1:100 and 1:1000 dilution, respectively. The samples were mounted in Mowiol-DABCO medium.

### STED nanoscopy and analysis

For STED nanoscopy, Leica TCS SP8 STED microscope with 100X, 1.4 NA PL APO oil objective was used to image growth cones. MTs labelled with Alexa Flour 568 were depleted with a continuous wave (CW) 660 nm depletion laser. While F-actin labelled with phalloidin-633 was depleted using a 775nm CW laser. Images were acquired by a HyD detector with time gating of 0.5 ≤ tg ≤ 6 ns with a pixel size of <36 nm. Images were deconvolved using the default settings and 15 iterations in the CMLE JM algorithm in the Huygens software (version 17.04.0p6 64b, Scientific Volume Imaging).

Neurons chosen for imaging were identified by the expression of a co-transfected soluble GFP plasmid construct. Any protrusion from the growth cone equal to or longer than 2µm was considered for filopodia analysis. The number and length of filopodia were measured manually using the line tool in the ImageJ software.

Analysis of alignment of MT to actin was done using the phallodin channel as reference to identify the initial position of alignment of MT with F-actin and the length of alignment was measured by tracking the MT along the actin bundle using the segmented line tool in ImageJ. For MT entry in filopodia a minimum incursion of 0.5 µm from the base of filopodia, was considered for analysis.

### Imaging and analysis for detyrosinated/total microtubules in the growth cone

For confocal imaging, Zeiss 780 LSM system was used with 63x/ 1.4 NA Plan-Apochromat oil objective. The images were acquired using a 405 nm diode laser (total MT), a 488 nm argon laser (detyrosinated MT or Fmn2) and a 561 nm DPSS laser (Alexa Fluor 568 phalloidin) for excitation with appropriate zoom to maintain pixel size of about ∼100nm.

For intensity measurements, imaging was conducted maintaining the same parameters across control and Fmn2-MO treated samples. The total MT image was thresholded (Otsu) using ImageJ and was used as a mask to measure the intensity of detyrosinated tubulin. The fluorescence was quantified using the formula, Corrected fluorescence = Integrated Density – (Area of selected cell x Mean background fluorescence). The detryosinated tubulin intensity was normalized to total tubulin intensity in the growth cone.

### Live-cell imaging of primary neurons

Growth cones were grown *in vitro* for 24 hours or 36 hours (rescue experiments). 5 µg of plasmid was used for transfections along with morpholino treatment. Only neurons with moderate expression levels were considered for imaging and analysis. Live-imaging of both EB3-GFP and EB3-mCherry (rescue experiments) were done on an IX81 system (Olympus Corporation) equipped with a Hamamatsu ORCA-R2 CCD camera using a 100X, NA 1.4 Plan Apo oil immersion objective. Wide-field imaging was done using the Xcellence RT (Olympus Corporation) software for 100 frames with a time interval of 2 seconds between each frame. During imaging the cultures were maintained at constant temperature of 37°C.

### EB3 comet tracking and data analysis

Comet tracking for the whole growth cone analysis was done using *plusTipTracker 1.1.4* version (Applegate et al., 2011, Matov et al., 2010) on MATLAB 2010b (Mathworks). The images were first background subtracted in ImageJ using a rolling ball radius of 50, to enhance the contrast. Detection of the comet was done via the plusTipGetTracks GUI with the following settings (Biswas & Kalil, 2018, Stout et al., 2014): detection method = anisotropic gaussian; PSF sigma = 1 to 1.5 and Alpha value= 0.005 to 0.02, depending on the most faithful detection of comets across several frames of each movie. Tracking of the comets were done using the following parameters: Search radius (Range) = 3 to 12; minimum sub-track length (frames) = 3; break non-linear tracks was unchecked; Maximum Gap Length (frames)= 11; Maximum shrinkage factor (relative to growth speed) = 0.8; Maximum angle (degrees) forward= 40 and backward = 20; fluctuation radius = 3.5. The post-processing parameters used were Frame rate (s) =2 and pixel size (nm)= 64. Raw values of growth speed, growth length, growth lifetime and dynamicity (collective displacement of all gap-containing tracks over the collective lifetimes) were exported to Graph Pad Prism 5 and used for graphical representation and analysis. Growth speed of comet tracks was visualized using the growth speed range from 0 to 20 µm/min in all frames using the *plusTipSeeTracks* GUI.

The EB3 comet tracking in the filopodia was conducted manually to avoid any false detection due to movement of the filopodia itself. Comets that emerged from the MT dense region were tracked using point tool of ImageJ, from the central domain to the peripheral or filopodial region, if they appeared in at least 3 continuous frames. The coordinates of the comet position were saved as ROI’s and exported to Microsoft Excel for calculating growth speeds. Comets that disappeared were observed for minimum 3-5 continuous frames to check for rescue. Only unambiguous comet entries into filopodia were considered for dwell time analysis. Comets entering the filopodia were manually tracked for growth speed and excursion depth analysis. Number of comets in each frame of the movie was extracted from the “movieinfo” file generated from *plusTipTracker* using a code written in MATLAB (Mathworks);

a=[100,1];

for i=1:100

a(i,1) =(length(movieInfo(i,1).xCoord));

end

The median value of comet numbers obtained from 100 frames of each movie was normalized to the area of the growth cone. The area of the growth cone was a mean of 10 measurements obtained manually using the freehand selection of ImageJ for every 10th frame.

### Protein purification

Actin was purified from rabbit muscle powder following standard protocols (Pollard, 1984) and labelled with Alexa 488 maleimide (Hansen et al., 2013). Porcine brain tubulin and rhodamine labelled tubulin was purchased from Cytoskeleton Inc. (Cat. # T240A) and (Cat. #TL590M-A), respectively. Bovine brain tubulin was used for dynamic co-localization studies of actin MT using TIRF microscopy. The GST-tagged chick Fmn2 constructs were expressed in BL21 Rosetta cells by inducing with 0.4 mM IPTG and purified following standard protocols (Harris & Higgs, 2006) using glutathione beads and further stored at 4°C in HEKG5 (20 mM HEPES, 1 mM EDTA, 50 mM KCl and 5% glycerol) buffer.

### Fmn2 interactions with F-actin or microtubules

Phalloidin stabilized F-actin was prepared from 30 μM G-actin by incubating actin with F-buffer (10 mM Tris pH 8.0, 0.2 mM CaCl2, 0.2 mM DTT, 2 mM MgCl2, 0.7 mM ATP, 50 mM KCl) for 1 hour at RT. 2 μM of Fmn2 FH2FSI and FH2_ΔFSI was incubated with increasing concentration of stabilized F-actin for another 30 mins.

Samples were centrifuged for 20 mins at 320,000 x g and analysed by SDS-PAGE. The amount of protein bound to actin in the pellet was quantified by densitometry on Commassie gels using ImageJ software (version 1.52g). The amount of protein bound to F-actin was plotted versus F-actin concentrations and the apparent dissociation constant was calculated by fitting a non-linear regression hyperbolic curve in Graph Pad Prism 5.0 software using the equation Y = Bmax * X / (Kd + X) where Bmax is maximum specific binding having same units as Y. Kd is denoted as the equilibrium binding constant, having same units as X and is the ligand concentration needed to achieve a half-maximum binding at equilibrium. In absence of filamentous actin, no protein was detected in the pellet which signified the solubility of the protein. To determine the affinity of MTs to Fmn2, 100 μM tubulin was polymerized at 37°C in BRB 80 (80 mM PIPES, 1 mM EGTA and 1 mM MgCl2) buffer with 1 mM GTP for 1 hour. 20 μM taxol was used to stabilize the MTs. 2 μM proteins were incubated with taxol stabilized MTs for another 1 hour in RT. Samples were centrifuged for 20 minutes at 100, 000 x g and analyzed by SDS-PAGE. The amount of protein bound to MTs was quantified as described above.

### Actin and microtubule bundling assays

Increasing concentrations of protein (FH2FSI and FH2_ΔFSI) was added to 2 μM preformed F-actin and allowed to form F-actin bundles in the presence of F-buffer. The amount of bundle formation was quantified after 30 minutes of incubation and centrifugation at 10,000 x g for 10 minutes to selectively pellet bundled F-actin. In case of F-actin control, the actin remained in the supernatant. The amount of actin fraction in the pellet and supernatant was quantified by densitometry on Coomassie-stained gels using the ImageJ software. The percentage of the actin pellet was calculated from the amount of actin in pellet divided by total amount of actin in the supernatant and pellet. The percentage of actin pellet was plotted against the increasing concentration of the protein using Graph Pad Prism 5.0 software.

In case of MT bundling assay, 2 μM of pre-formed MTs were incubated with increasing concentrations of the protein (FH2FSI and FH2_ΔFSI) for 30 minutes and centrifuged at 5,000 x g for 10 minutes. The amount of MT fraction in the pellet and supernatant was quantified by densitometry on Coomassie-stained gels using the ImageJ software. Further, the percentage of MT in the pellet was calculated and plotted in a similar manner as described above for actin pellets.

### Microtubule stabilization assay

MTs were polymerized from 100 µM tubulin in presence of BRB 80 buffer (80 mM PIPES, 1mM EGTA and 1 mM MgCl2) with 1 mM GTP for 1 hour. 20 µM Taxol was used to stabilize the MTs. Increasing concentrations of FH2FSI was incubated with the MTs for 30 min at RT. To assess the MTs stability to cold-induced depolymerisation in presence of FH2FSI, the samples containing MTs and FH2FSI and their respective control samples were incubated on ice for another 30 min and spun in a bench-top ultracentrifuge (TLA 100.3; Beckman Coulter) for 20 mins at 100,000g at 4°C to isolate cold-stable MTs. Supernatant and pellets were resolved on 10% SDS PAGE and the proteins were detected by Coomassie staining.

### Cross-linking of F-Actin and microtubules

2 µM FH2FSI or 2 μM FH2_ΔFSI constructs were incubated with 2 μM pre-formed phalloidin-stabilized actin and 2 μM taxol-stabilized MT in the presence of BRB 80 buffer for 30 minutes and centrifuged at 2,000 x g for 10 minutes. Supernatants and pellets were analyzed by SDS-PAGE. The amount of F-actin and MTs in the pellet was quantified individually following the same method as described earlier.

### TIRF microscopy and evaluation of concomitant microtubule and actin assembly

Perfusion chambers were prepared using glass slides and cover-slips using double-sided sticky tape and were coated with silane PEG (Creative works) following standard protocols (Portran et al., 2013). Alexa-488 phalloidin-labelled F-actin, in absence or presence of either FH2FSI or FH2_ΔFSI, was imaged using the inverted microscope (ApoN/TIRF 100x /1.49 oil immersion objective on an IX 81 system; Olympus) equipped with the Cell TIRF module (Olympus) and an Hamamatsu ORCA-R2 CCD camera in TIRF buffer (0.2% methylcellulose, 3 mg/ml glucose, 20 µg/ml catalase and 100 µg/ml glucose oxidase). To observe the MT bundles under TIRF microscope, 5% rhodamine labelled tubulin and unlabeled tubulin was used to form taxol stabilized MTs and incubated with the protein (either FH2FSI or FH2_ΔFSI). Co-alignment of preformed F-actin and taxol-stabilized MT, in presence or absence of either FH2FSI or FH2_ΔFSI, was also imaged similarly using the TIRF mode of Olympus IX 81 microscope.

Actin and MT growth inside the perfusion chamber were initiated by flowing 1 µm actin monomers (12% Alexa-488 labelled G-actin), 25 µM tubulin dimers (10% ATTO-565 labelled tubulin) and labelled MT seeds in the absence or presence of 0.5 μM FH2FSI or FH2_ΔFSI. The buffer used for the reaction contains 10 mM HEPES, 50 mM KCl, 5 mM MgCl2, 1 mM EGTA, 20 mM DTT, 1% BSA, 0.1 mM ATP along with the TIRF buffer. Samples were visualized using a TIRF microscope (Eclipse Ti, Nikon with an iLAS TIRF system) equipped with a EMCCD camera (Evolve 512, Photometrics) and controlled by the Metamorph software (Molecular Devices). The actin and MT samples were observed using an Apochromat 100x/1.49 oil immersion objective. Simultaneous dual color time-lapse imaging was performed at 35°C for actin/tubulin co-assembly using a dual emission splitter. Images were acquired for 12 minutes at 1 frame per 20 secs with 100 ms exposure time. All image analysis was conducted using the ImageJ software and a custom written co-alignment analysis macro (Prezel et al., 2017).

### Microcontact printing and substrate-choice border assay

PDMS stamps containing zig-zag pattern of 100 or 150 µm width stripes and 10 µm height were made from a silicon master (custom manufactured by Bonda Technology Pte. Ltd., Singapore). The silicon wafer was silanized with trichloromethylsilane vapours in vacuum for 5 minutes. After keeping the master in the vacuum desiccator overnight, the process was completed by heating the master plate at 70⁰C for 30 minutes. A 10:1 mixture of PDMS and curing agent was degassed by centrifugation at 3000 rpm for 5 minutes. The mixture was poured on the silicon master and cured at 60⁰C for 2 hours. Microcontact printing on glass coverslips was done using protocol as adapted from (Thery & Piel, 2009). The stamps were inked with 100 µl solution of tetramethylrhodamine (TMR) -BSA (2.5 µg/ml) + laminin (10 µg/ml) + Fibronectin (10 µg/ml) for 20 minutes at RT. After removing the excess solution, the stamps were washed with 1x PBS and allowed to dry for a minute before printing onto the PLL-coated (0.1 mg/ml) glass coverslips. PBS was added to prevent drying of the pattern on the 35 mm glass-bottomed dish.

Alternating borders of poly-L-lysine and ECM proteins were created by the microcontact printing method described above. Neurons co-transfected with a soluble GFP-expressing plasmid along with control or Fmn2 morpholinos were grown on the patterned substrates for 36 hours before imaging. Only neurons, whose cell bodies were on the laminin + fibronectin patterns (red stripes) and axons that approached the borders were considered for analysis. Imaging was done using a 40x/1.4 NA oil objective on an Olympus IX81 microscope and dual-channel (GFP and TMR) images were captured. Data were pooled from four independent biological replicates.

### *In vivo* imaging and analysis of EB3 comets in zebrafish embryos

All procedures employed were approved by the Institutional Animal Ethics Committee (IAEC) of IISER Pune, India. Wild-type TU strain zebrafish maintained at 28.5 ℃ under a 14 h light and 10 h dark cycle, housed in a recirculating aquarium (Techniplast) were crossed to obtain eggs. The eggs were co-injected with 25 pg Ngn1: EB3-GFP, 25 pg Ngn1: TagRFP-CAAX (both gifts from Dr Mary Halloran, University of Wisconsin, Madison, USA) and 0.125 mM morpholino (Gene Tools, LLC) in a 2 nL volume in the cytoplasm at 1 cell stage. The embryos were maintained at 28.5 ℃ in E3 medium (5 mM NaCl, 0.17 mM KCl, 0.33 mM MgSO4, 0.33 mM CaCl2) containing 0.002% methylene blue until the time of imaging.

For depleting Fmn2 levels *in vivo*, the splice-blocking Fmn2 morpholino (Zf_Fmn2-MO) used was 5’-ACAGAAGCGGTCATTACTTTTTGGT -3’ and the standard control morpholino(Ctl-MO) used was 5’-CCTCTTACCTCAGTTACAATTTATA -3’. Zf_Fmn2-MO is expected to block the splicing of the intron between exons 5 and 6, resulting in the introduction of an early stop codon and an increase in size of the mRNA. The efficacy of the Zf_Fmn2-MO was evaluated using RT-PCR (Figure S2B). The injected embryos were screened for mosaic expression of GFP and RFP, dechorionated and mounted laterally. Vacuum grease (Dow Corning) was used to make a well on the glass slide wherein the zebrafish embryo was positioned laterally in E3 medium containing 2% methylcellulose (Sigma-Aldrich) and 0.03% MS-222 (Sigma-Aldrich) and sealed with a cover slip. Growth cones of Rohon-Beard (RB) sensory neurons labelled by the Ngn promoter driven EB3-GFP and Tag-RFP in 24-28 hpf embryos were imaged on a Zeiss LSM 780 confocal microscope using 63x/1.4 NA oil immersion objective for 100 frames every 1.5 or 2 second intervals.

For analysis of EB3 comets obtained from *in vivo* imaging in zebrafish, kymographs were generated using the segmented line tool and KymographBuilder in ImageJ and velocities and lifetimes were extracted using the velocity measurement tool.

### Graphical representation and statistical analysis

All the graphical representation and statistical analyses were performed using GraphPad Prism 5. In text representation of values is mean ± standard error of mean (SEM). Graphical representations are scatter dot plots with longer middle line indicating the mean and error bars indicating the SEM. For box plots, the box extends from the 25^th^ to 75^th^ percentiles, the line inside the box is plotted at the median and the whiskers represent the smallest and largest values. Number of data points quantified in each graph has been indicated either in the relevant text or figure legends. Mann-Whitney U test was employed for comparing two groups. When comparing more than two groups, non-parametric one-way ANOVA or Kruskal-Wallis test was used followed by Dunn’s multiple comparison post-test.

## ETHICS APPROVAL

All protocols used in this study were approved by Institutional Animal Ethics Committee and the Institutional Biosafety Committee of IISER Pune.

## AVAILABILITY OF DATA AND MATERIAL

All data generated or analysed during this study are included in this published article. The raw data are available from the corresponding author on reasonable request.

## AUTHORS’ CONTRIBUTIONS

Conceptualization: T.K., P.D. and A.G.; Investigation and analysis: T.K. (all experiments involving primary cultures of neurons, imaging & analysis of live zebrafish growth cones), P.D. (all *in vitro* and biochemical experimentation), D.N. (zebrafish experimentation); Writing of manuscript: T.K., P.D., D.N. and A.G.; Reagents and protocols: S.M. (preparation of rabbit muscle actin and advice on biochemistry protocols); Funding Acquisition: A.G.. All authors gave final approval for publication and agree to be held accountable for the work performed therein.

## ACKNOWLEDGMENTS

The authors are grateful to Dr. Laurent Blanchoin (CytomorphoLab, Université Grenoble-Alpes/CEA/CNRS/INRA, Grenoble, France) for hosting P.D. and providing technical expertise in establishing the TIRF-based actin and microtubule co-polymerization assays. Jérémie Gaillard and Christophe Gurein in the Blanchoin laboratory are acknowledged for providing purified bovine tubulin and rabbit skeletal muscle actin. The authors thank Dr. G. Deshpande, Princeton University, Dr. L. Blanchoin, Université Grenoble-Alpes/CEA/CNRS/INRA, Grenoble, Dr. S. Rath, IISER Pune and Dr, N. K. Subhedar, IISER Pune for their critical reading of this manuscript. Preliminary biochemical experiments were conducted with goat tubulin provided by Kunalika Jain & Dr. Chaitanya Athale, IISER Pune. The authors acknowledge the IISER Pune Microscopy Facility and the National Facility for Gene Function in Health and Disease (NFGFHD) at IISER Pune for access to equipment and infrastructure.

## Supplementary Information

**Fig. S1.**
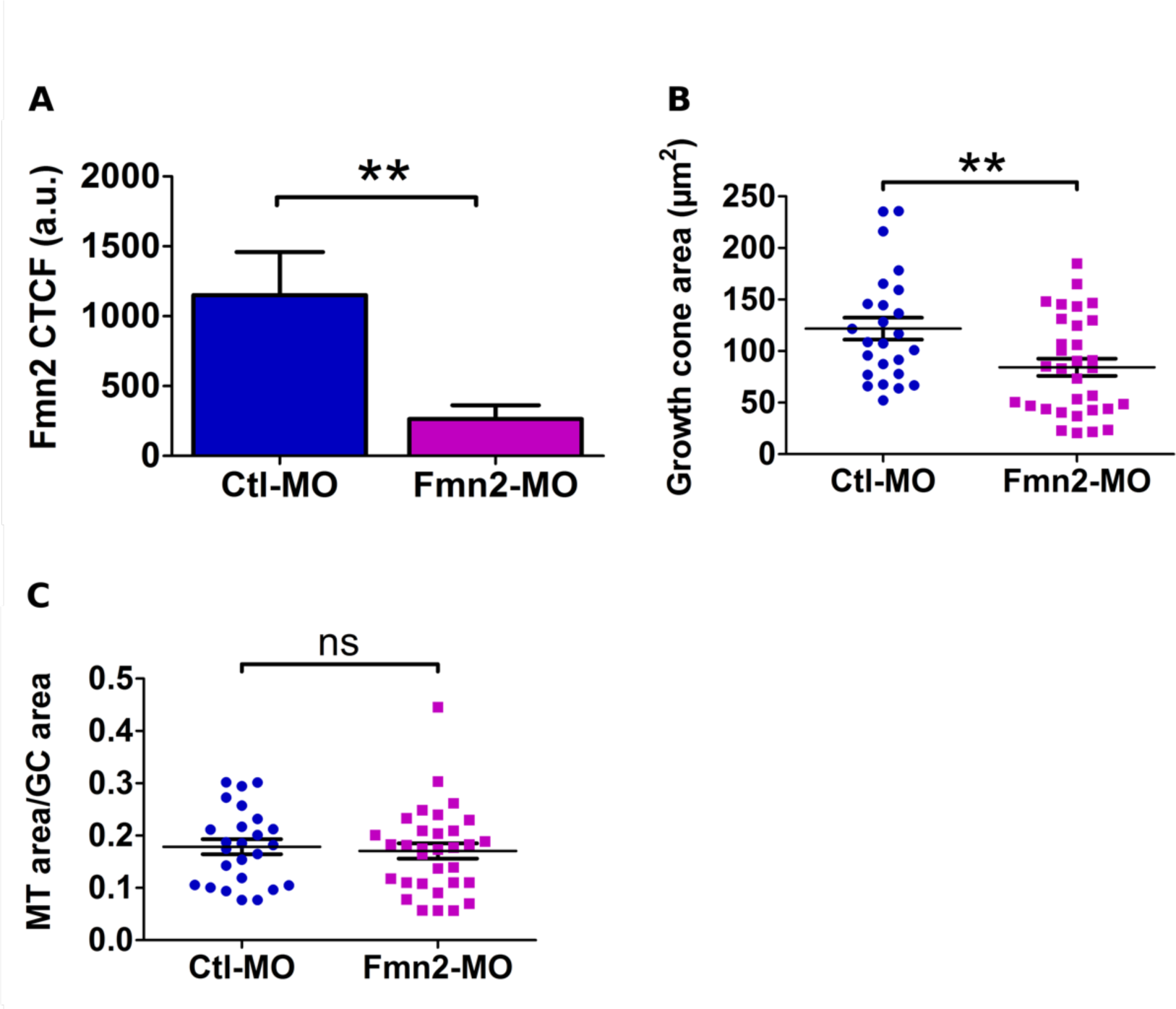
Fmn2 knockdown affects growth cone area. (A) Quantification of endogenous levels of Fmn2 in Ctl-MO (n=13) and Fmn2-MO (n=10) transfected growth cones of chick spinal neurons. Corrected total cell fluorescence (CTCF) = Integrated density – (Area of the growth cone x mean background intensity). a.u., arbitray units. (B) Quantification of growth cone area and (C) area occupied by MT in the growth cone normalized to growth cone area. The area occupied by microtubules is identified using anti-alpha tubulin antibody and calculated by intensity thresholding within the area outlined as the growth cone. (Ctl-MO, n=25; Fmn2-MO, n=32 for both B & C). **P<0.01, ns P>0.05; Mann-Whitney test.

**Fig. S2.**
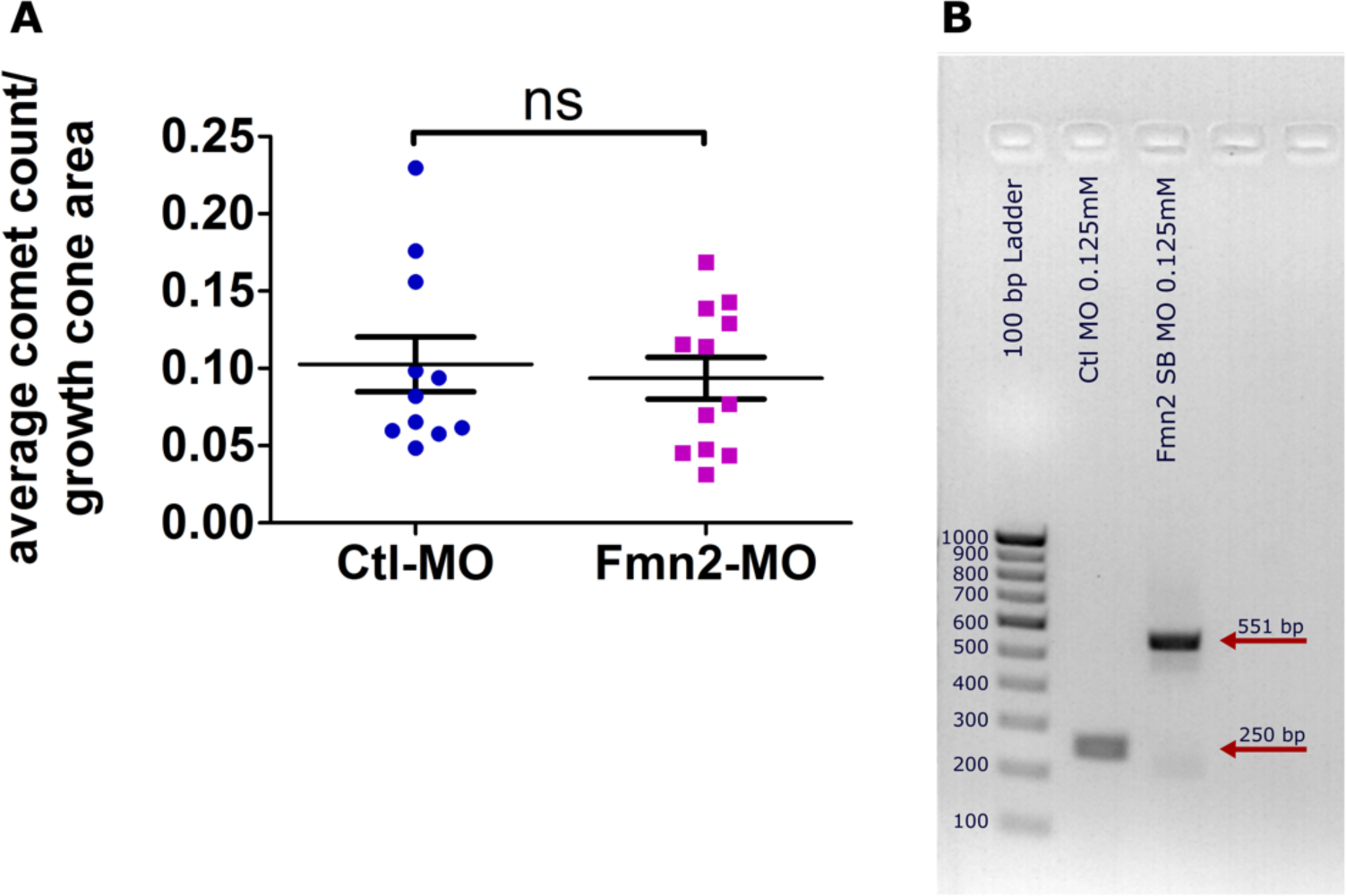
EB3 comets in Ctl-MO and Fmn2-MO growth cones and efficacy of the Zf_Fmn2-MO. (A) Quantification of average comet count normalized to growth cone area for Ctl-MO (n= 11) and Fmn2-MO (n=12). ns P>0.05; Mann-Whitney test. (B) RT-PCR with specific primers in exon 5 and 6 of zebrafish Fmn2 shows that the Zf_Fmn2-MO efficiently blocks removal of the intron between exons 5 and 6 leading to a larger amplicon (Lane 3, 551 bp) as compared to control (Lane 2, 250 bp).

**Fig. S3.**
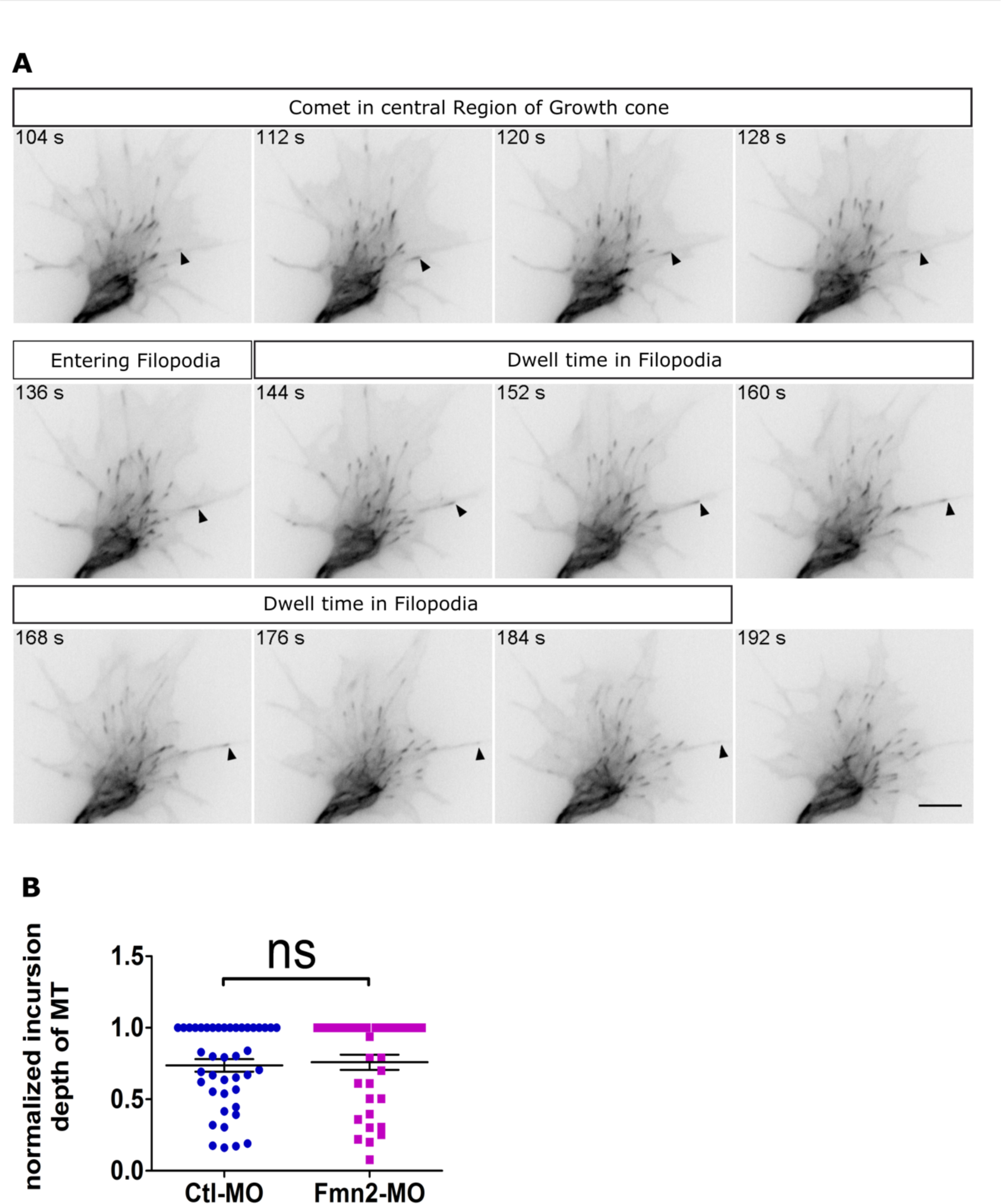
EB3 comets in filopodia. (A) Representative snapshots of a timelapse series showing EB3 dynamics in a growth cone. The arrowhead marks a EB3 comet entering the filopodia from the central region of the growth cone and its dwell time in the filopodia before it disappears. (B) Quantification of the distance covered by an individual EB3 comet inside filopodia normalized to total filopodial length in Ctl-MO (n= 42) and Fmn2-MO (n= 35) growth cones. ns P>0.05; Mann-Whitney test.

**Fig. S4.**
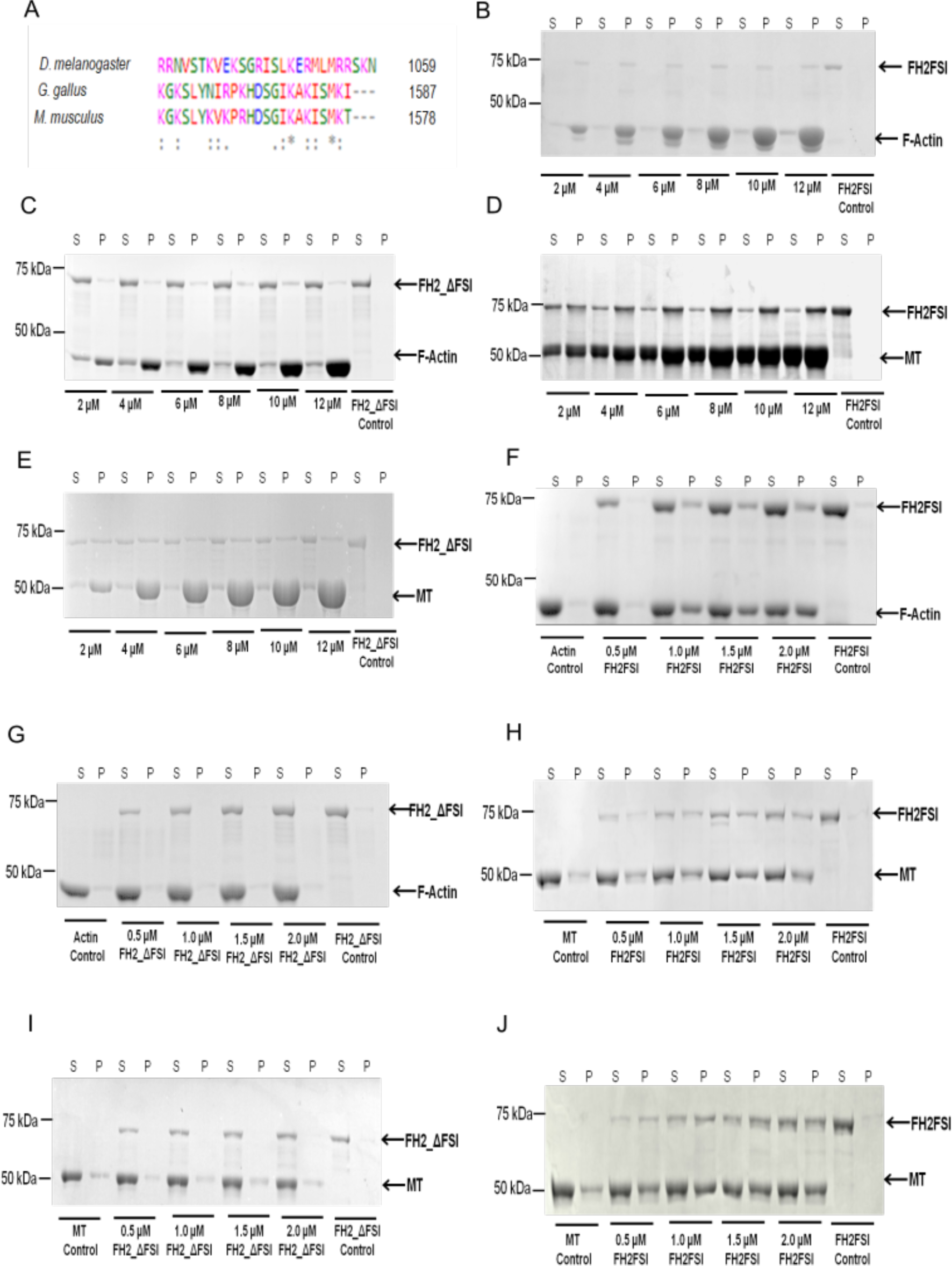
Interaction of Fmn2 with F-actin and Microtubules. (A) Alignment of FSI domain protein sequences (UniProt Protein Database) for *D. melanogaster* (Q24120), *G. gallus* (A0A1B2VWD2) and *M. musculus* (Q9JL04) using Clustal Omega (1). The colour coding (Magenta: basic amino acids, Green: uncharged polar amino acids, Red: non polar hydrophobic amino acids, Blue: Acidic amino acids) and the symbols (*conserved residues, : strongly similar residues, · weakly similar residue) are used according to the Clustal Omega. (B) High-Speed (320,000 x g) co-sedimentation assay of increasing concentration of F-actin with FH2FSI shows direct interaction between FH2FSI and F-actin. (C) High speed (320,000 x g) co-sedimentation assay of increasing concentration of F-actin with FH2_ΔFSI showing reduced association. (D) High speed (100,000 x g) co-sedimentation assay of MT with FH2FSI shows direct interaction of MT with FH2FSI. (E) High speed (100,000 x g) co-sedimentation assay of FH2_ΔFSI with increasing concentration of MT indicates reduced association. (F) Low speed (10,000 x g) co-sedimentation assay with F-actin indicates bundling of actin filaments in the pellet fraction by FH2FSI. (G) Low speed (10,000 x g) co-sedimentation assay of FH2_ΔFSI with F-actin shows compromised F-actin bundling activity. (H) Bundling of MTs in the pellet fraction in presence of FH2FSI in low speed (5,000 x g) co-sedimentation assay. (I) Reduced MT bundles in the pellet in the presence of FH2_ΔFSI in a low speed (5,000xg) co-sedimentation assay. (J) FH2FSI stabilizes MTs against cold induced depolymerization. Representative SDS-PAGE showing self-assembled MTs are resistance to cold induced depolymerisation when incubated with FH2FSI. After centrifugation at (100,000xg), MTs can be visualized only in the supernatant in the control. With the increasing concentration of FH2FSI, MTs can be visualized in the pellet. Note: In each gel image “S” represents the supernatant fraction and “P” represents the pellet fraction.

**Fig. S5.**
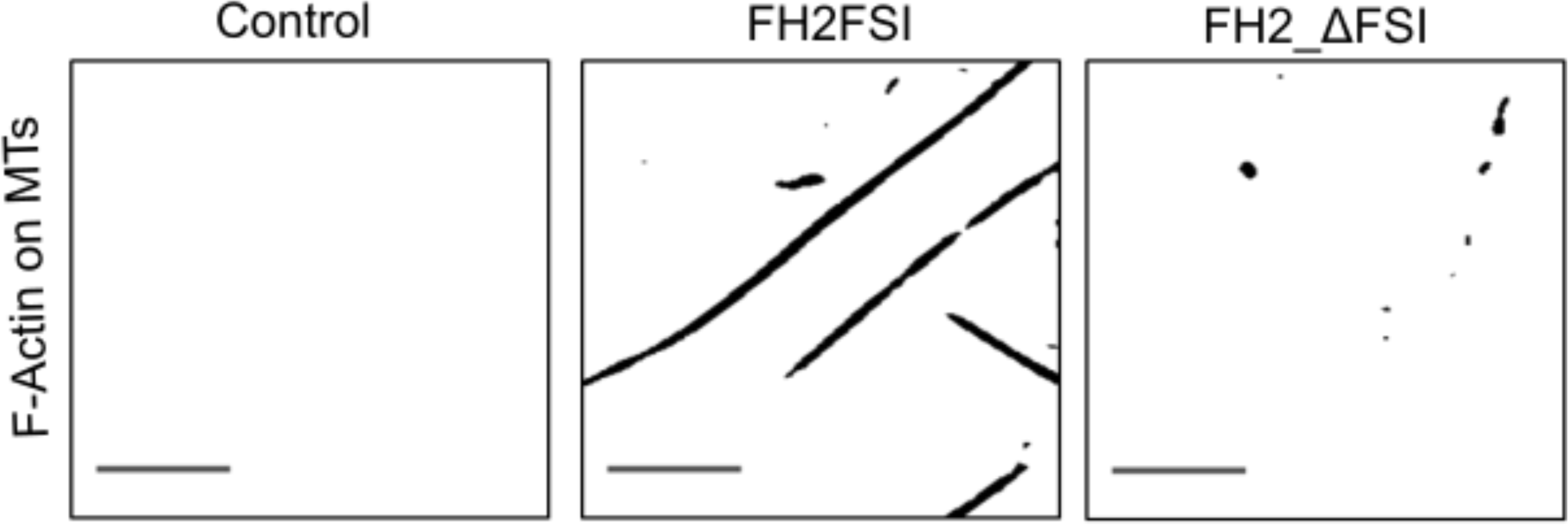
Fmn2 cross-bridges polymerizing Microtubules and F-actin filaments. The masked images represent the actin area that co-localizes with microtubule in the last frame of the movies in Control, FH2FSI and FH2_ΔFSI respectively. Scale bar 10 µm.

**Table S1.**
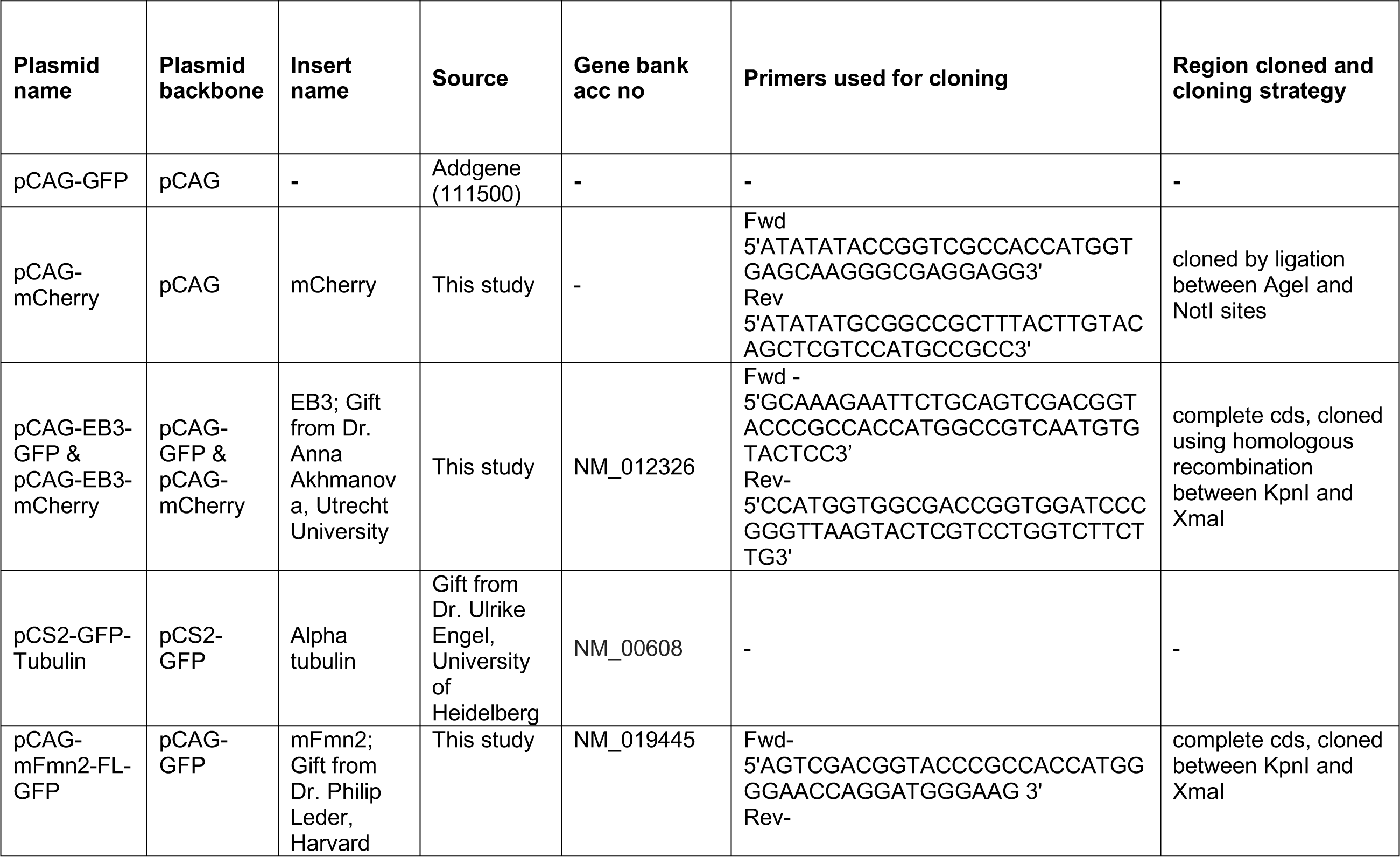
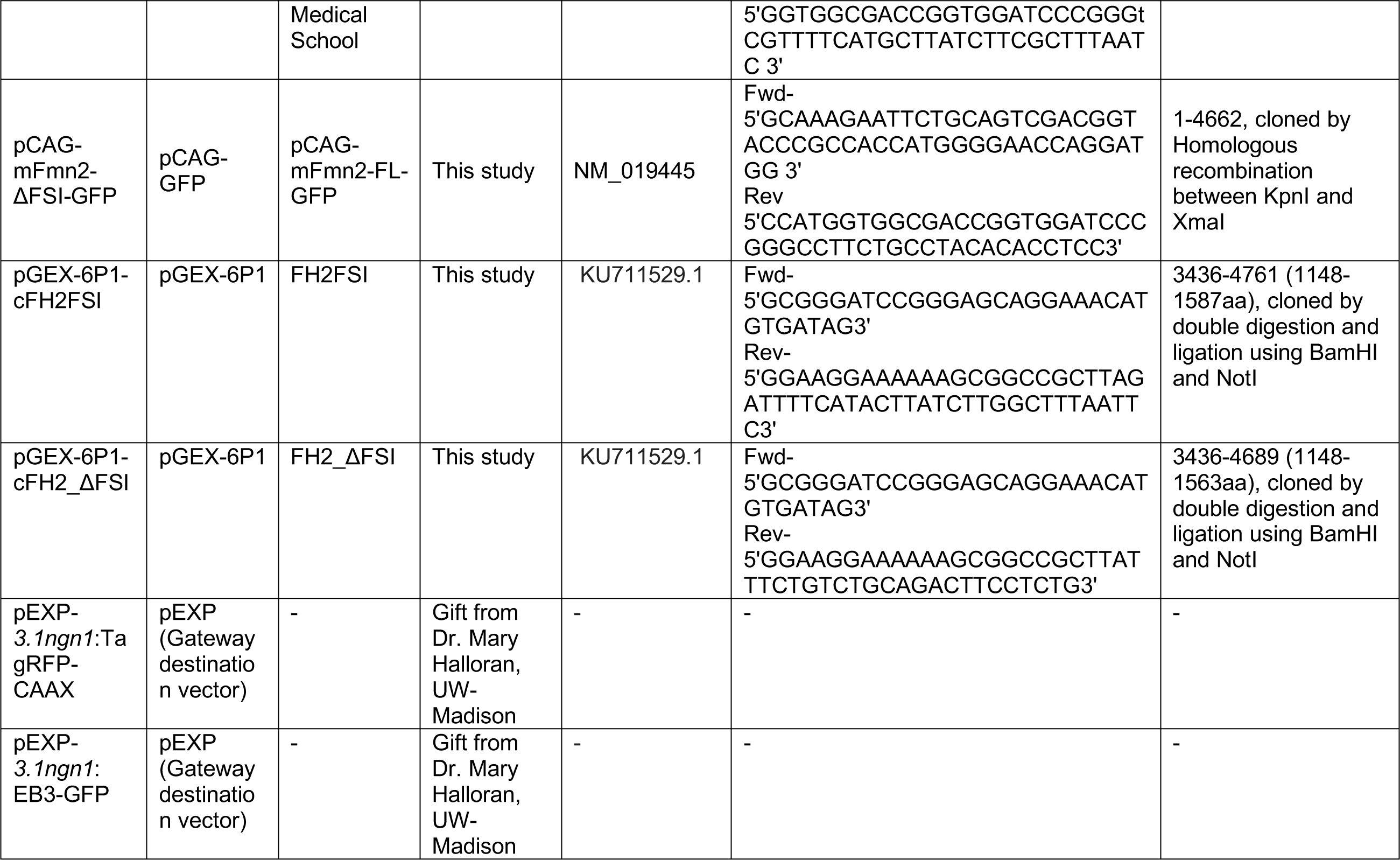
Plasmid constructs used in this study indicating source, vector backbone, cloning strategy and primers used for cloning.

**Movie S1 (separate file). Ctrl-MO transfected growth cone displaying EB3-GFP comets.**

The EB3 comets are color-coded in correspondence with increasing growth speed (blue to red) using the plustiptracker software.

**Movie S2 (separate file). Fmn2-MO transfected growth cone displaying EB3-GFP comets.**

The EB3 comets are color-coded in correspondence with increasing growth speed (blue to red) using the plustiptracker software.

**Movie S3 (separate file). RB neuron growth cone of a Ctl-MO injected zebrafish displaying EB3 comets *in-vivo*.**

**Movie S4 (separate file). RB neuron growth cone of a Zf_Fmn2-MO injected zebrafish displaying EB3 comets *in-vivo*.**

**Movie S5 (separate file). Co-polymerization of actin and tubulin in the absence of protein.**

Dual colour TIRFM timelapse video of actin (green) and tubulin (magenta), polymerizing together in absence of protein. The time (in secs) represents the elapsed time start from the acquisition. The scale bar is 10 μm.

**Movie 6 (separate file). Co-polymerization actin and tubulin in the presence of FH2FSI.**

Dual colour TIRFM timelapse video of FH2FSI induced co-assembly of actin (green) and tubulin (magenta). The time (in secs) represents the elapsed time start from the acquisition. The scale bar is 10 μm.

**Movie 7 (separate file). Co-polymerization of actin and tubulin in the presence of FH2_ΔFSI.**

